# Genetic Variation in Photosynthetic Maintenance and Post-Chill Recovery for Early Season Maize Plantings

**DOI:** 10.64898/2026.10.30.763178

**Authors:** Tyler J McCubbin, Emma N Leary, Donald Coon, Jacob D Washburn

## Abstract

Earlier maize planting could extend the growing season and improve synchronization of crop nitrogen demand with biogenically available soil nitrogen in spring, but seedling photosynthesis and development are strongly constrained by chilling. Eleven genetically diverse maize inbred lines were evaluated under a spring-like controlled environment regime containing five days of chilling stress followed by a recovery period, and across four early-spring field environments. Gas exchange, chlorophyll fluorescence, chlorophyll status, immature-leaf growth, vegetative stage, biomass, and visible cold injury were quantified to characterize physiological and developmental responses during chilling and recovery. Chilling substantially reduced net CO₂ assimilation and rapidly decreased ΦPSII, while stomatal conductance responded slower. Genotypic differences in assimilation and ΦPSII were compressed during prolonged chilling but reemerged during recovery. Chilling also slowed immature-leaf elongation and largely arrested vegetative developmental progression; both resumed rapidly after warming, and advanced recovery stage development was associated with a greater fraction of mature leaf biomass. Assimilation and ΦPSII were most tightly associated during recovery and remained positively associated across field environments, although the strength of this relationship depended strongly on environment. Field expression of genotype effects was highly environment dependent, but broad-sense heritability was moderate to high for a variety of measured photosynthetic and physiological traits. Compared with the benchmark chilling-tolerant C_4_ grass *Miscanthus,* the maize panel exhibited similar photosynthetic performance, underscoring the accession dependence of this comparative benchmark. Taken together, these results identify photosynthetic maintenance and recovery as separable components of early-season cold performance and characterize physiologically informative and heritable traits associated with chilling stress resilience.

## Introduction

C_4_ photosynthesis supports some of the world’s most productive crops, including maize (*Zea mays* L.), sorghum (*Sorghum bicolor*), and sugarcane (*Saccharum officinarum*). C_4_ lineages exhibit considerable biochemical and evolutionary diversity (Washburn *et al*., 2015, 2021), and provide particularly high photosynthetic efficiency under conditions of high light and warm temperature. However, because of the system’s tropical ancestry advantages are less pronounced at low temperature maize production in temperate regions (Ehleringer and Björkman, 1977; Zhu *et al*., 2008; Long and Spence, 2013). Maize growth and photosynthesis are strongly inhibited by suboptimal temperature, with chilling injury occurring as temperatures approach or fall below approximately 10°C (Greaves, 1996; Long and Spence, 2013). This cold sensitivity limits the extent to which maize can exploit the early spring growing season. The Circular Economy that Reimagines Corn Agriculture (CERCA) initiative has proposed that increasing seedling cold tolerance could enable earlier planting, extend the annual period of C_4_ carbon assimilation, and better synchronize maize growth and nitrogen demand with the spring period of natural soil nitrogen availability (Ojeda-Rivera *et al*., 2025). Accordingly, development of maize capable of maintaining growth and photosynthetic function under early-season chilling conditions represents an important component of the proposed CERCA ideotype.

Photosynthesis is among the most chilling-sensitive physiological processes in maize (Sowiński *et al*., 2005; Hund *et al*., 2008). Following the transition to autotrophic growth, continued seedling development depends on the capacity to capture energy from light and maintain carbon assimilation despite the limitations imposed by suboptimal temperatures to chloroplast development, C_4_ metabolism, and the photosynthetic electron transport system (Greaves, 1996; Long and Spence, 2013). Chilling decreases the abundance or activity of important photosynthetic enzymes, including pyruvate phosphate dikinase (PPDK), phosphoenolpyruvate carboxylase (PEPC), and RubisCO, while simultaneously increasing susceptibility of the photosynthetic apparatus to excess excitation energy (Naidu *et al*., 2003; Long and Spence, 2013). These effects are particularly important when low temperature occurs under high irradiance, when limitations to carbon metabolism can increase excitation pressure on photosystem II (PSII) and contribute to photoinhibition (Pimentel *et al*., 2005; Grzybowski *et al*., 2019). Chlorophyll fluorescence therefore provides a complementary measure of photosynthetic function to gas exchange measurements. In particular, the operating quantum efficiency of photosystem II (ΦPSII) reflects the proportion of absorbed light being used in PSII photochemistry under light-adapted conditions and provides a measure of the capacity for photosynthetic electron transport (Murchie and Lawson, 2013). Maintenance of carbon assimilation during chilling may therefore depend not on a single physiological process, but on coordination among carbon fixation, PSII photochemistry, stomatal behavior, and protection or recovery of the photosynthetic apparatus. Moreover, these photosynthetic traits also exhibit useful genetic variation. Chlorophyll fluorescence has long been proposed as a selection tool for cold tolerance of photosynthesis in maize (Fracheboud *et al*., 1999), and more recent work has identified genetic variation affecting PSII efficiency during early growth under cold conditions (Urzinger *et al*., 2025).

The strong chilling sensitivity of maize appears to be an avoidable consequence of C_4_ photosynthesis. Several species within the genus *Miscanthus*, a member of the Andropogoneae tribe and relatively close to maize and other major C_4_ grasses, are notable for their ability to remain photosynthetically productive at low temperature (Beale and Long, 1995; Naidu *et al*., 2003; Dohleman *et al*., 2009; Long and Spence, 2013). Comparisons between maize and *Miscanthus × giganteus* have shown substantially greater maintenance of CO_2_ assimilation in *Miscanthus* during chilling. Whereas maize grown at low temperature can exhibit reductions in photosynthesis approaching 80%, *M. × giganteus* maintains a much greater proportion of its warm-grown photosynthetic capacity (Naidu *et al*., 2003). This response has been associated with maintenance of C_4_-cycle and Calvin-cycle enzymes, including PPDK and RubisCO, as well as rapid repair of PSII and increased photoprotective capacity (Naidu *et al*., 2003; Long and Spence, 2013; Spence *et al*., 2014). These observations demonstrate that sustained C_4_ photosynthesis at low temperatures is physiologically possible and provides a useful comparative model for improvement of maize. However, chilling tolerance also varies within *Miscanthus*, and much of the mechanistic understanding of cold-tolerant C_4_ photosynthesis has been derived from a limited number of exceptionally tolerant genotypes (Głowacka *et al*., 2015; Ojeda-Rivera *et al*., 2025). Thus, *Miscanthus* provides an informative benchmark for chilling-tolerant C_4_ photosynthesis but may not necessarily define a uniform species-level response against which maize can be compared (Głowacka *et al*., 2014).

Following domestication from teosinte in southwestern Mexico, maize diversified through farmer selection and adaptation as it spread throughout the Americas and subsequently into temperate production regions worldwide (Vigouroux *et al*., 2008; Hufford *et al*., 2012; Flint-Garcia, 2013; Larkins, 2017). This history generated extensive phenotypic and genetic diversity among landraces and modern germplasm, including adaptation to high elevations, northern latitudes, and cooler growing environments. Highland maize and Northern European germplasm provide prominent examples of adaptation to cooler environments (Eagles and Lothrop, 1994; Rodríguez *et al*., 2010), and genetic variation for seedling cold tolerance has been identified in diverse maize panels (Revilla Temiño *et al*., 2014; Yi *et al*., 2021). Multiple loci and natural variants associated with low-temperature responses have subsequently been identified (Revilla Temiño *et al*., 2014; Zeng *et al*., 2021; Jiang *et al*., 2022), demonstrating that cold tolerance is amenable to genetic improvement.

Despite this progress, the physiological architecture underlying genetic variation in maize chilling tolerance remains partially resolved. Many studies have focused on germination, emergence, survival, or individual photosynthetic traits, while fewer have examined how carbon assimilation, PSII photochemistry, stomatal behavior, chlorophyll status, and growth respond together across genetically diverse germplasm during both chilling and subsequent recovery. Recovery is particularly important because genetic differences that are not apparent during the period of maximal stress may be expressed after favorable temperatures return. For example, several European Flint genotypes exhibit strongly reduced leaf growth during chilling but have been shown to rapidly resume growth after returning to a warm temperature; this response was described as a physiological “standby mode” (Riva-Roveda *et al*., 2016). However, this and many other detailed physiological studies have evaluated relatively small sets of maize genotypes, often under light environments substantially below the 1,000-2,000 µmol m^−2^ sec^−1^ encountered during clear spring days in the U.S. Corn Belt. Despite the genetic variation of cold tolerance being well established, it remains unclear how much capacity for maintenance of photosynthesis under chilling and high irradiance that maize germplasm possesses. In addition, the physiological processes that govern that capacity exhibit genetic variation that can be exploited through breeding.

The present study addressed the question: What capacity does maize germplasm have for maintenance of photosynthesis during chilling stress, and what physiological processes are essential to chilling and recovery? A genetically diverse panel of 11 maize inbred lines was evaluated under controlled chilling and recovery conditions designed to approximate physiologically relevant early-season cold stress while maintaining comparatively high irradiance. Gas exchange, PSII photochemistry, chlorophyll status, immature-leaf growth, developmental progression and biomass accumulation were quantified to characterize variation in photosynthetic performance and its relationship to whole plant growth and recovery from chilling stress. *Miscanthus* accessions were included as comparative C_4_ germplasm representing a lineage in which exceptional chilling tolerance of photosynthesis has previously been documented. The controlled-environment analyses were complemented by field experiments testing whether physiological and growth responses translated across naturally variable early-season cold environments. Finally, broad-sense heritability was estimated for component photosynthetic and growth traits to assess the extent of genetic control and their potential utility as breeding targets in the development of an earlier-planted, cold-tolerant maize ideotype. Together, these experiments link photosynthetic maintenance and recovery with vegetative development, field performance, and the trait architecture relevant to development of the cold-tolerant CERCA ideotype.

## Materials and Methods

### Plant material and controlled-environment growth

Eleven maize inbred lines were selected to span broad germplasm and heterotic backgrounds: 3IIH6 (Iodent); B73 and LH244 (Stiff Stalk); B97, PHK76, and PHN46 (Non-Stiff Stalk); CML247, CML322, and NC358 (Tropical); Il14H (Sweet Corn); and F2 (European Flint). Seed was obtained through the USDA National Plant Germplasm System. The panel included public and ex-PVP material and was used consistently across controlled-environment and field analyses. Two *Miscanthus* accessions, Pi417947 and a *Miscanthus × giganteus* hybrid, were obtained from the USDA GRIN stock center and from Gilbert H. Wild and Son, LLC (Reeds, MO, USA).

Seeds were sown in 3-inch-deep, 36-cell Star Plug trays containing a 3:1 mixture of Pro-Mix BX (Premier Tech LTD., Rivière-du-Loup, QC, Canada) and Turface MVP (Profile Products, Buffalo Grove, IL, USA). After 10-14 d, three to five seedlings per genotype were transplanted into 1.72-gallon (6.5 l) nursery pots (A.M.A. Horticulture, BM600, Kingsville, ON, Canada) filled with Pro-Mix BX amended with 2 Tsp (10.2 ml) of a blend consisting of one part Osmocote Plus (ICL Growing Solutions, Geldermalsen, Netherlands) 15-9-12 slow-release fertilizer, one part Plant Doctor Tomato Maker (Organic Laboratories, Inc., Stuart, FL, USA) 4-2-6 cal mag, 1/12 part Sprint 330 (BASF, Ludwigshafen am Rhein, Germany), and 1/100 part ferrous sulfate (Diamond Brand, Cary, NC, USA). Pots were wrapped with R-3 reflective insulation, the soil surface was covered with an insulated disk, and pots were placed on R-10 rigid foam board on top of a short bench.

Plants were grown in Conviron walk-in chambers (Controlled Environments Limited, Winnipeg, MB, Canada). The prechill and recovery program was designed to approximate early-season Corn Belt conditions associated with substantially earlier planting. Early May, when approximately half of the Illinois maize crop is typically planted (Irwin and Good, 2012), was used as an agronomic reference for the warm-stage environment. A 22/16 °C day/night regime was selected for prechill and recovery, with the overnight temperature chosen to remain above temperatures that commonly impair normal maize development (Miedema, 1982; Riva-Roveda *et al*., 2016). The intervening chilling treatment used 12/10 °C day/night temperatures, near the lower thermal boundary for maize vegetative growth (Ojeda-Rivera *et al*., 2025). Plants experienced 5 days (D) of prechill, 5 D of chilling, and 5 D of recovery under a 14-h photoperiod with photosynthetic photon flux density ramped from 300 to a peak of 1,200 μmol m⁻² s⁻¹, for a daily light integral of 51.7 mol photons m⁻² day⁻¹. Relative humidity was programmed to 60% during Prechill, 40% during Chilling, and 60% during Recovery stages. To prevent the occurrence of thermal gradients, two horizontal fans were suspended from the ceiling 1m above canopy height, while one fan was placed on the opposite side of the chamber on the floor, circulating air under the table. Air at canopy level and soil temperatures were verified using EasyLog EL-SIE-2+ loggers (Lascar Electronics, Erie, PA, USA) and CR1000X dataloggers (Campbell Scientific, Logan, UT, USA), respectively.

### Sampling design and physiological measurements

Measurements were organized around the transition into and out of chilling; the controlled environment experiment was replicated four times. Gas exchange and light-adapted fluorescence were collected at the end (D5) of the Prechill stage, at the beginning and end (D1 and D5) of the Chilling and Recovery stages. Growth and SPAD measurements were collected at D1 and D5 of each stage, allowing growth to be estimated directly within each five-day interval, and vegetative stage was recorded at the D5 endpoint of each stage. The youngest actively growing leaf was reassessed at the beginning of each stage; consequently, immature-leaf growth is interpreted as a within-stage D1-to-D5 response rather than as a continuous trajectory of the same organ across all three stages.

Gas exchange and chlorophyll fluorescence were measured with an LI-6800 Portable Photosynthesis System equipped with a Multiphase Flash Fluorometer and 2-cm² chamber (LI-COR Environmental, Lincoln, NE, USA). Measurements were made on the youngest fully expanded mature leaf. Chamber irradiance was set to 1,200 μmol m⁻² s⁻¹ for light-adapted measurements, from which net CO₂ assimilation, stomatal conductance, ΦPSII, and light-adapted Fv′/Fm′ were obtained. Leaf temperature in the LI-6800 chamber was controlled to match the stage environment (22 °C during prechill and recovery; 12 °C during chilling). Fluorescence nomenclature and interpretation followed established practice (Murchie and Lawson, 2013). Dedicated dark-adapted fluorescence was measured on the same relevant leaf after at least 1 h of dark adaptation. For paired same-leaf measurements, NPQ was calculated from dedicated dark-adapted Fm and the corresponding light-adapted Fm′ as (Fm − Fm′)/Fm′.

For morphology, the mature source leaf and the youngest actively growing immature leaf were measured for length and width. Mature-leaf relative chlorophyll content was measured with a Konica Minolta SPAD-502DL meter (Konica Minolta, Tokyo, Japan); five readings distributed across the illuminated leaf surface were averaged for each leaf. Developmental stage was assigned from the number of fully mature leaves; a leaf was considered mature only when the ligule was fully formed (Hanway, 1966). At final harvest, plant material was separated into immature leaves, mature leaves, stem, and roots and dried at 65 °C for at least 4 d before dry-mass determination.

### Controlled-environment measurements and statistical analysis

Quality control for LI-6800 measurements was adjudicated on a trait-specific basis so that a technical problem with one measurement channel did not automatically invalidate other measurements from the same plant and timepoint. Inclusion criteria for gas-exchange and simultaneous light-adapted fluorescence measurements were based on whether leaf temperature and measurement conditions matched the intended experimental stage; instrument identity was retained as quality-control metadata. Technically valid negative assimilation values were retained, whereas negative stomatal-conductance values and failed all-zero fluorescence measurements were treated as invalid. Dark-adapted fluorescence was evaluated independently of the gas-exchange temperature filter removing any need for quality control selection.

Longitudinal analyses of assimilation, stomatal conductance, light-adapted F_V_′/F_M_′, and ΦPSII used mixed models containing genotype × timepoint and experimental-run × timepoint effects with repeated measurements on individual plants. Timepoint-specific residual variances were allowed; ΦPSII was analyzed on the logit scale and displayed on the response scale, whereas the other primary physiology traits were analyzed on their observed scales. Genotype estimated marginal means and Tukey-adjusted comparisons were used for stage-specific statistical inference. Statistical model selection for all photosynthesis traits is described in Supplementary Table S1. Vegetative stage at Prechill D5, Chilling D5, and Recovery D5 was analyzed as a repeated numeric response with genotype × timepoint and experimental-rep × timepoint effects and a plant random intercept. Mature-leaf SPAD was analyzed separately using the controlled-environment experiment with complete six-timepoint coverage, with D5 analyses across additional experiment replicates used as sensitivity checks. Dark-adapted F_V_/F_M_ was analyzed cross-sectionally at D5. Relationships among assimilation, ΦPSII, dark-adapted Fv/Fm, growth, SPAD, developmental stage, and endpoint biomass allocation were evaluated using linear models with experimental run included where supported. Biomass responses were analyzed at the individual-plant level, and allocation was evaluated using both tissue ratios and allometric comparisons among biomass components. All statistical analysis for controlled environment and field experiments was performed using R language programming (R Core Team, 2026).

### Early-spring field experiments

The maize panel was evaluated in four early-spring field environments over 2025 and 2026 at two locations: Location A refers to the University of Missouri Rollins Bottoms Experiment Station (38.92 N, −92.35W; Haymond silt loam soil) while Location B refers to the University of Missouri Genetics Farm (38.90 N, −92.27W, Mexico silt loam soil). V3 seedlings were transplanted into three replicate blocks at each site, with five plants per genotype per block targeted at establishment. A cold sensitive Dekalb commercial hybrid (P1213AM; LibertyLink®/Roundup Ready® 2) was included as a field phenotyping reference but was excluded from the genomic and heritability analysis. Plants were germinated and grown before transplant in the same way as described above for the controlled environment experiments. Upon transplant, they were exposed to ambient field conditions for 5 d. Air temperature, 4-inch soil temperature, PPFD, relative humidity, and precipitation were recorded for each environment (https://agebb.missouri.edu/). Visible cold injury was scored daily and summarized as the fraction of plants within each genotype × block that remained free of visible injury.

Immature-leaf length and mature-leaf SPAD were measured immediately before transplanting and again after the five-day field exposure. Immature-leaf growth was calculated over this interval, and mature-leaf SPAD at the endpoint was also expressed relative to the pretransplant value. Net CO₂ assimilation, stomatal conductance, ΦPSII, and light-adapted F_V_′/F_M_′ were collected at the field endpoint. When multiple plants were measured within a genotype × environment × block, plant subsamples were averaged to a single block-level experimental unit before statistical analysis.

### Field statistical analysis

Genotype effects were evaluated separately within each field environment using mixed models with genotype as a fixed effect and block as a random effect. Tukey-adjusted genotype comparisons were used when the omnibus genotype effect was significant. A pooled analysis across all four environments evaluated genotype × year × location effects with block nested within an environment. Trait-relationship figures used the same block-level data nested within environment. The assimilation–ΦPSII relationship was summarized across all environments by linear regression; assimilation–immature-leaf growth and stomatal conductance–SPAD relationships were evaluated as complementary field associations.

### Genomic marker validation and population structure

Marker calls shared between the maize HapMap and Genomes to Fields (G2F) resources were evaluated for cross-source concordance. Seven highly concordant accessions selected from 79 accessions represented in both resources were used as trusted controls. Shared SNPs were assigned stability tiers from full-panel discordance and leave-one-accession-out validation; the highest-confidence Platinum tier required complete concordance across the trusted set and validation in every leave-one-out analysis. The final panel contained 1,310 SNPs distributed across all 10 maize chromosomes. These markers were used to construct the realized genomic relationship matrix for the 11 study lines and to summarize population structure by principal-component analysis. The workflow and raw data are hosted electronically.

### Broad-sense heritability across field environments

Broad-sense heritability was estimated from the field block means for the 11 maize inbred lines across four environments. The multi-environment phenotype model included environment as a fixed effect and genotype, genotype × environment, and block-within-environment as random effects. This phenotype-only best linear unbiased predictor (BLUP) model, which assumes independent genotype effects, was treated as the primary analysis. A complementary genomic BLUP (GBLUP) model replaced the identity covariance among genotype effects with the realized genomic relationship matrix from the SNP panel. Entry-mean H², Cullis H², and the mean pairwise difference reliability (mean ΔH²) were calculated for the phenotype model as described (Schmidt *et al*., 2019b,a); genomic Cullis and mean ΔH² estimates accounted for the realized genetic variance of each genotype pair. Stomatal conductance was excluded *a priori* because block-level coverage was insufficient for reliable multi-environment variance estimation.

## Results

### A modified growth chamber system imposed repeatable spring-like chilling events

A repeatable combination of high light intensity and low temperatures was considered critical for testing hypotheses about early-season photosynthesis in the U.S. Cornbelt region. However, achieving these conditions in growth chambers is technically challenging due to factors including heat generation from high intensity lamps, thermal gradients, higher heat exchange of potting conditions compared to field soils, and the increasing difficulty of humidity control as temperatures decrease. Considerable time and experimentation were invested to develop systems for buffering these issues and monitoring protocols to ensure conditions were maintained. Supplementary Fig. S1 demonstrates the success of the growth chamber modifications and configurations in maintaining desired diurnal temperatures (both soil and air), soil moisture, and humidity levels during each phase of the experiment and across experiment repetitions. The resulting schema therefore imposed a defined transient cold event within an otherwise spring-like growth regime, without imposing frost damage. Of the 165 plants grown across the trials 96% were measured at all timepoints in Prechill, Chilling, and Recovery stages.

### Chilling rapidly inhibits carbon assimilation and PSII photochemistry

Net CO₂ assimilation, stomatal conductance, ΦPSII, and light-adapted F_V_′/F_M_′ were followed from prechill through chilling and recovery to determine how maize photosynthesis responds to sustained chilling (Fig. 1). Mean assimilation across the entire maize panel declined from approximately 8.4 μmol CO₂ m⁻² s⁻¹ at Prechill D5 to 3.3 at Chilling D1 and 1.8 at Chilling D5, corresponding to an approximately 78% reduction over the chilling period. Stomatal conductance increased slightly at Chilling D1 before returning near the prechill value by Chilling D5. In contrast, ΦPSII fell sharply from approximately 0.163 before chilling to 0.055 at Chilling D1, increasing slightly to 0.090 by D5 of the Chilling period. Light-adapted F_V_′/F_M_′ declined more gradually from approximately 0.259 before chilling to 0.201 by Chilling D5. Thus, the rapid loss of carbon assimilation at the onset of chilling coincided more closely with reduced PSII operating efficiency than with stomatal closure.

**Figure 1.**
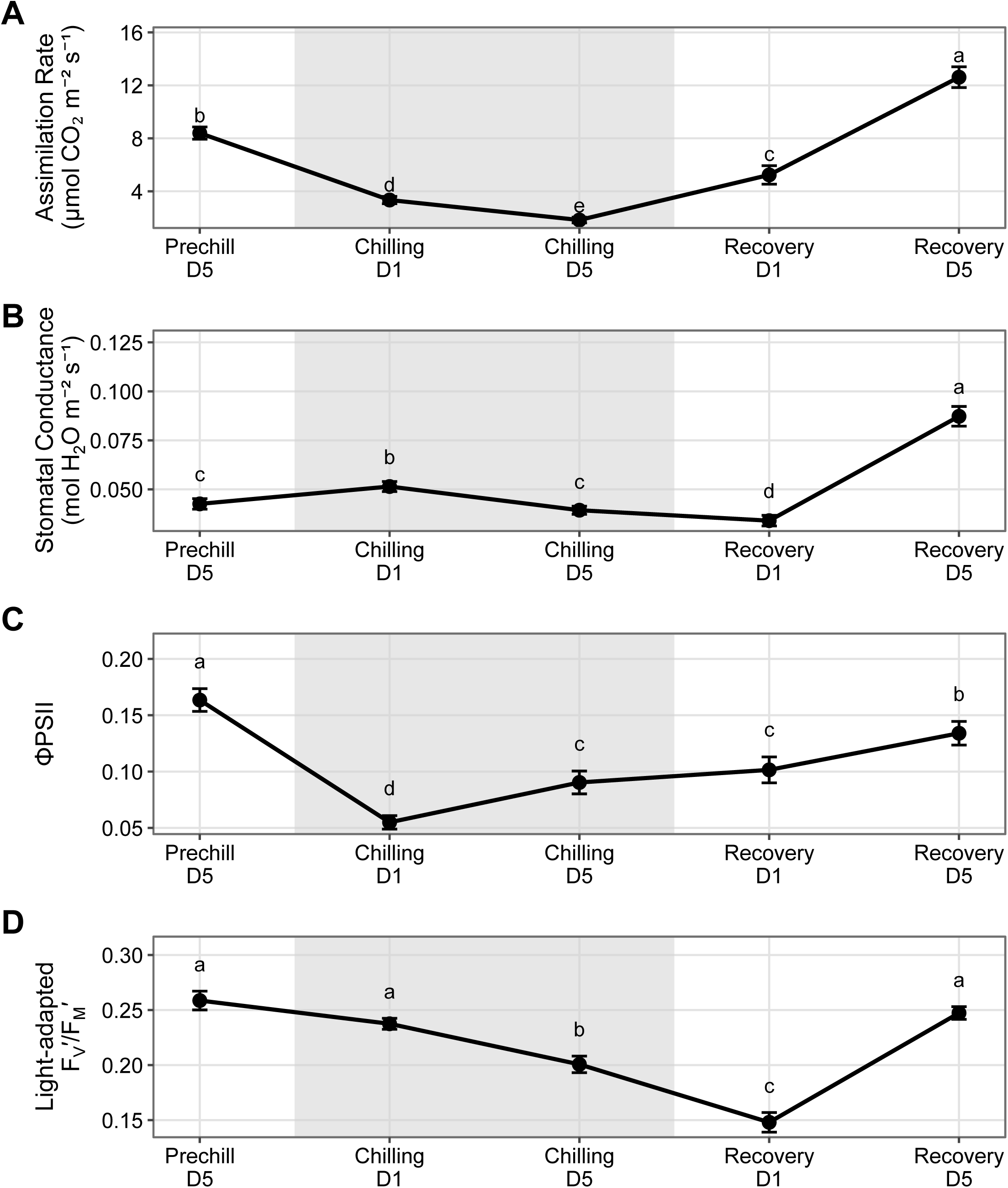
Photosynthetic physiology of the maize diversity panel across prechilling, chilling, and recovery. (A) Net CO₂ assimilation, (B) stomatal conductance, (C) photosystem II quantum yield (ΦPSII), and (D) light-adapted F_V_′/F_M_′ at the indicated experimental timepoints. The shaded interval denotes the five-day chilling treatment. Points represent the mean of model-adjusted genotype estimated marginal means (EMMs) at each timepoint, and error bars represent the SE among the 11 genotype EMMs. Statistical analyses included 703, 678, 529, and 539 valid plant-time observations for assimilation, stomatal conductance, ΦPSII, and light-adapted Fv′/Fm′, respectively. Different letters indicate Tukey-adjusted differences among timepoints at P < 0.05. Longitudinal inference used mixed models containing genotype × timepoint and experiment × timepoint effects, a plant random intercept, and timepoint-specific residual variances; ΦPSII was modeled on the log scale, untransformed values displayed.

Photosynthetic recovery began immediately after the return to spring-like temperatures in the recovery stage. Assimilation increased to approximately 5.2 μmol CO₂ m⁻² s⁻¹ at Recovery D1 and 12.6 by Recovery D5, while stomatal conductance rose to approximately 0.087 mol H₂O m⁻² s⁻¹. Recovery of fluorescence traits followed a different trajectory: light-adapted Fv′/Fm′ reached its lowest mean value at Recovery D1 before returning near the prechill value by Recovery D5, and ΦPSII recovered more gradually to approximately 0.134. Carbon gain therefore recovered rapidly after chilling even while PSII fluorescence parameters continued to readjust during the early recovery period.

Two *Miscanthus* accessions were included as physiological comparisons for C_4_ chilling performance in three growth chamber experiments (Supplementary Fig. S4). During both chilling timepoints, neither accession maintained higher assimilation than the maize panel, and ΦPSII converged to similarly low values across maize and *Miscanthus*. Differences re-emerged during recovery, but the direction and magnitude of those differences were not consistent across the two accessions. These data show that the expected superior maintenance of chilling photosynthesis was not uniformly expressed in the *Miscanthus* material evaluated here.

Photosynthetic trajectories varied among maize genotypes, with significant genotype × timepoint interactions for assimilation, stomatal conductance, light-adapted F_V_′/F_M_′, and ΦPSII. Stage-end estimates illustrate the strongest feature of these interactions: assimilation was distinct among genotypes before chilling, but no pairwise differences remained at Chilling D5. Substantial differentiation reappeared by Recovery D5 (Fig. 2A). ΦPSII followed the same pattern of a chilling compression, but genotype-specific responses were only seen during recovery (Fig. 2B).

**Figure 2.**
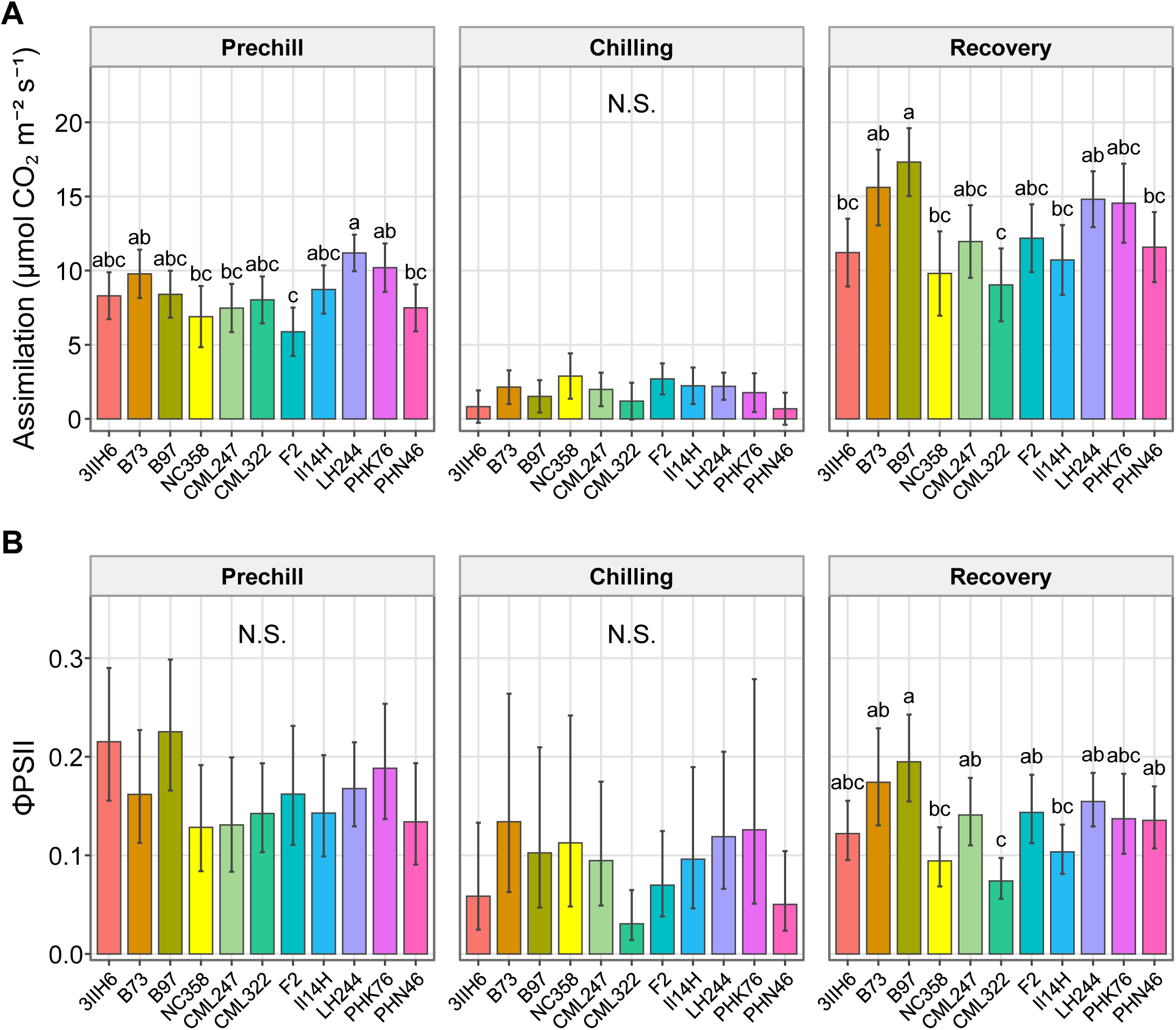
Genotypic variation in photosynthetic traits at the end of each experimental stage. Genotype-specific EMMs for (A) net CO₂ assimilation and (B) ΦPSII at Prechill D5, Chilling D5, and Recovery D5 for the 11 maize inbred lines. Error bars represent 95% confidence intervals around the EMMs; ΦPSII estimates are displayed on the response scale following analysis on the logit scale. The longitudinal datasets contained 703 valid plant-time observations for assimilation and 529 for ΦPSII. Within each stage, genotypes not sharing a letter differed at Tukey-adjusted *P* < 0.05. EMMs and contrasts were obtained from longitudinal mixed models containing genotype × timepoint and experiment × timepoint effects, a plant random intercept, and timepoint-specific residual variances.

Several genotypes exhibited distinct stage-dependent responses. LH244 exhibited the highest assimilation before chilling, whereas B97 displayed the strongest photosynthetic recovery. At Recovery D5, B97 had the highest assimilation (17.3 μmol CO₂ m⁻² s⁻¹), stomatal conductance (0.124 mol H₂O m⁻² s⁻¹), light-adapted F_V_′/F_M_′ (0.291), and ΦPSII (0.195) among the maize genotypes. Prolonged chilling therefore compressed expression of genetic differences in photosynthetic performance, while return to favorable temperature revealed substantial genetic variation in recovery capacity.

### Coordination between PSII operating efficiency and carbon assimilation changes through chilling and recovery

Because carbon assimilation depends on continued conversion of absorbed light into biochemical energy, the relationship between assimilation and ΦPSII was evaluated at the D5 endpoint of each stage with differences among controlled-environment experiment reps considered. Assimilation and ΦPSII were moderately associated before chilling and strongly associated during recovery., Conversely, the relationship during chilling was weak and the observed range of both traits was compressed (Fig. 3). For all genotypes, a steep positive association was generally evident during recovery, while chilling values fell within a narrow range of both low-assimilation and low-ΦPSII (Supplementary Fig. S5). These data show that operating PSII efficiency tracked carbon assimilation most closely during recovery, when variation in both traits expanded beyond the narrow variability observed in the chilling stage.

**Figure 3.**
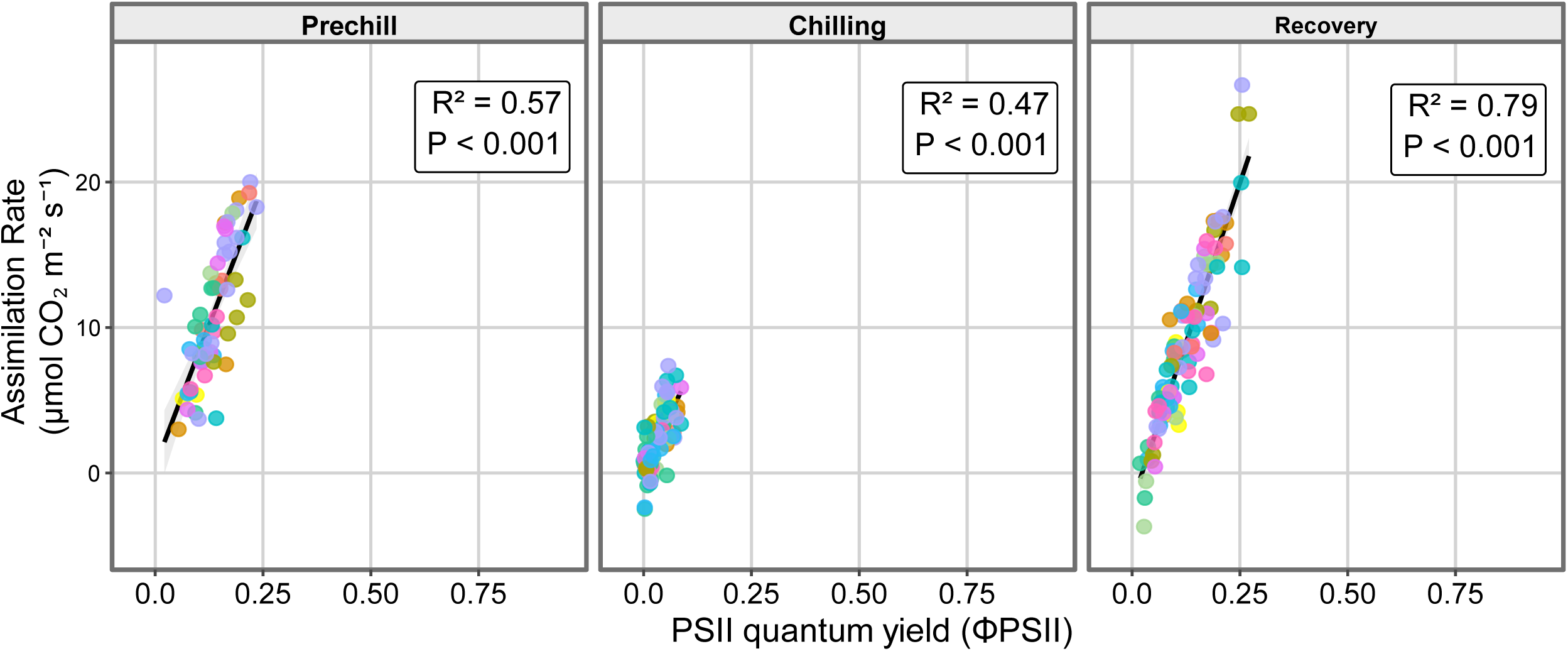
Relationship between net CO₂ assimilation and ΦPSII at the end of prechilling, chilling, and recovery. Net CO₂ assimilation is plotted against ΦPSII for individual maize plants at Prechill D5, Chilling D5, and Recovery D5, with points colored by genotype. Black lines show the linear relationship within each stage, with shaded 95% confidence intervals. Panel annotations report the R2 for ΦPSII after adjustment for Experiment and the corresponding P-value. Analyses included *N* = 59, 88 and 99 observations for Prechill, Chilling, and Recovery, respectively (N = 246 total).

Dark-adapted F_V_/F_M_ provided a complementary measure of PSII status. Mean dark-adapted F_V_/F_M_ declined from approximately 0.72 before chilling to 0.50 at Chilling D5 and recovered to approximately 0.73 by Recovery D5 (Supplementary Fig. S6). Genotypic differences were detectable for Prechill D5 but were not resolved at Chilling D5 or Recovery D5. Nonphotochemical quenching (NPQ) showed no detectable genotypic differentiation at the D5 endpoint of any experimental stage, while mean NPQ was greater during chilling and recovery than before chilling (Supplementary Fig. S7A). Relationships of NPQ with assimilation rate and mature-leaf SPAD were stage dependent, with no clear resolvable relationships (Supplementary Fig. S7B,C). Thus, variation in operating PSII efficiency was more closely aligned with variation in carbon assimilation than were dark-adapted maximum PSII efficiency or NPQ.

### Chilling restricts immature-leaf growth and vegetative developmental progression

The same actively growing immature leaf was marked at D1 and measured again at D5 within each stage to determine whether inhibition of photosynthetic function was accompanied by reduced vegetative growth and development. Vegetative stage was also recorded from fully collared leaves at the D5 endpoint of each stage (Fig. 4A). Mean rate of immature-leaf elongation declined from 22.5 mm d^−1^ at Prechill to 14.8 mm d^−1^ during Chilling and recovered to 158% of the Prechill amount by Recovery D5 (35.6 mm d^−1^). Genotypes differed in growth before chilling, whereas genotype differences were not detected during the chilling or recovery intervals. Chilling therefore restricted expansion of developing leaf tissue while compressing detectable genetic variation in this short-term growth response.

**Figure 4.**
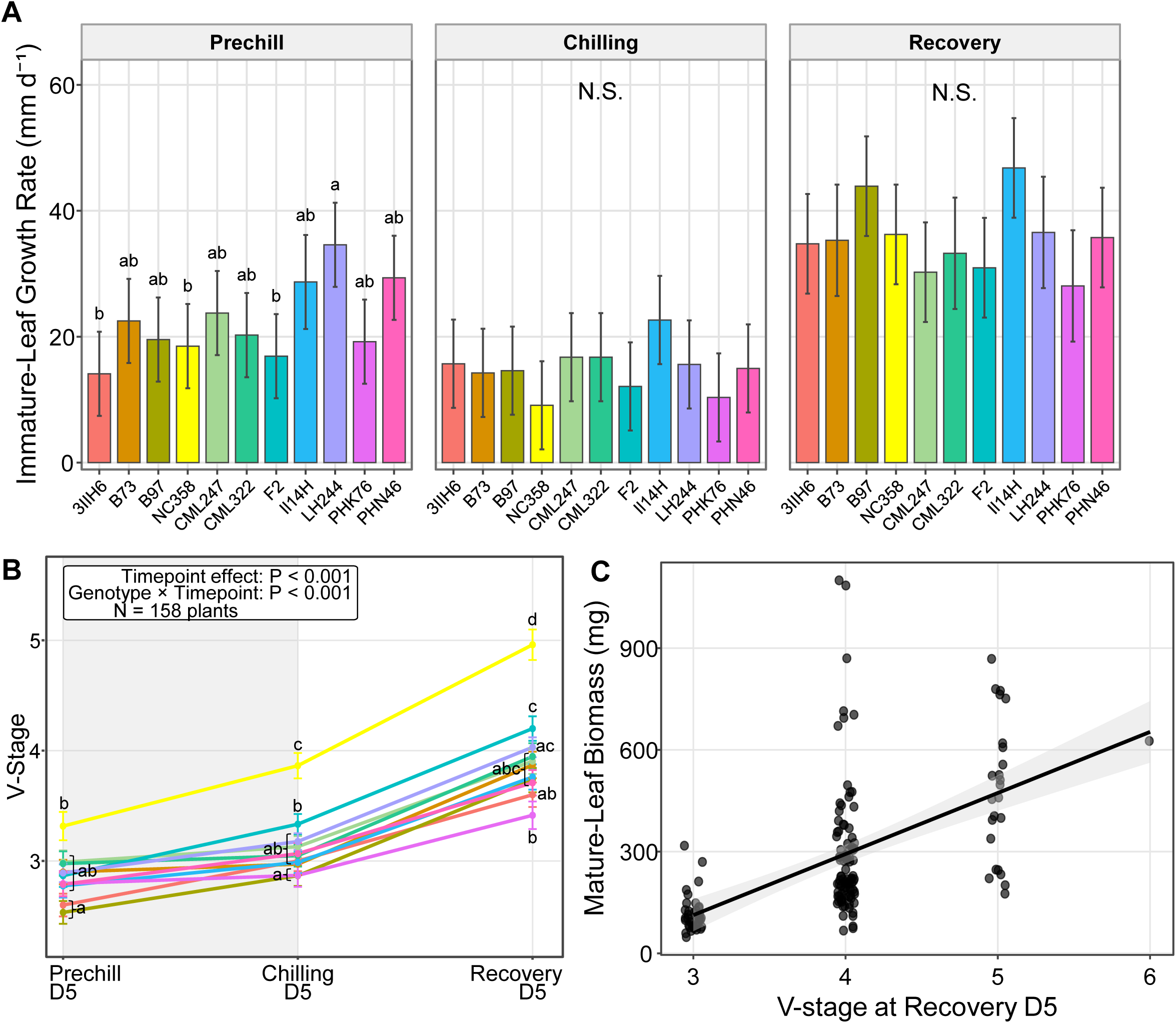
Vegetative development and biomass allocation during chilling and recovery. A) Immature-leaf growth rate for the 11 maize genotypes across all experiment stages, calculated from repeated measurements of the marked immature leaf within each stage. (B) D5 vegetative stage across Prechill, Chilling, and Recovery. (C) Mature-leaf biomass fraction is plotted against Recovery-D5 V-stage using matched endpoint biomass observations; the line shows the experiment-adjusted linear trend, shaded region shows 95% CI. N = 54, 55 and 51 for prechill, chilling and recovery stages in (A); 158, and 191 for B and C, respectively. Within a stage or biomass trait, genotypes not sharing a letter differed at Tukey-adjusted P < 0.05.

Developmental progression showed a similar response; mean vegetative stage increased only from V2.86 to V3.12 during the five-day chilling interval but reached V3.92 after five days of recovery. Most plants failed to advance a complete vegetative stage during chilling, whereas most developed by at least one stage during recovery. Genotype-specific trajectories differed across the treatment, with the strongest separation evident by Recovery D5 (Fig. 4B). Thus, chilling delayed both expansion of the actively growing leaf and the emergence of successive fully mature leaves, whereas recovery restored both elongation and developmental progression.

Genotypic differences were evident for immature leaf, root, and shoot biomass, whereas no statistically significant differences were observed among genotypes for mature leaf biomass. Partitioning between above- and belowground biomass diverged among genotypes, with the shoot:root ratio ranging from approximately 1.13 in PHK76 to 1.88 in F2 (Supplementary Fig. S8). Vegetative stage at Recovery D5 was positively associated with the fraction of leaf biomass present as mature tissue (Fig. 4C). Thus, developmental progression during recovery was reflected as a higher fraction of mature leaf biomass than developing tissue at the conclusion of the experiment.

For the majority of genotypes assessed, mature-leaf SPAD increased across the prechill period, declined during chilling, and recovered after restoration of warm conditions (Supplementary Fig. S9A). SPAD trajectories differed among genotypes through most of the six-timepoint sequence, although genotype differences were no longer detected at Recovery D5. Chlorophyll status therefore retained a substantial genotype component while following a pattern distinct from the rapid changes in carbon assimilation and ΦPSII. Mature-leaf SPAD was positively associated with stomatal conductance at the D5 endpoint of Prechill, Chilling, and Recovery, although substantial variation remained within each stage (Supplementary Fig. S9B).

### Carbon assimilation remains coupled to ΦPSII in the early-spring field experiments, but the strength of association is environment-dependent

The same maize panel was evaluated in a natural experiment consisting of four early-spring field environments spanning two locations and two years. Air and soil temperature, PPFD, humidity, and precipitation differed substantially among the four five-day exposure periods, but approximately reflect the environment imposed in the controlled environment experiments with more extreme overnight low temperatures for three of the four experiments (Fig. 5A). The increased chill resulted in distinct patterns of cold injury which could be visually scored (Fig. 5B). Injury developed most rapidly and extensively in the 2025 Location A environment, whereas cold resistance remained comparatively high across most genotypes in 2026 Location A; the two Location B environments produced intermediate but genotype-dependent injury trajectories (Fig. 5C). The field experiments therefore extended the controlled-environment analysis into naturally variable combinations of low temperature, high irradiance, humidity, and precipitation.

**Figure 5.**
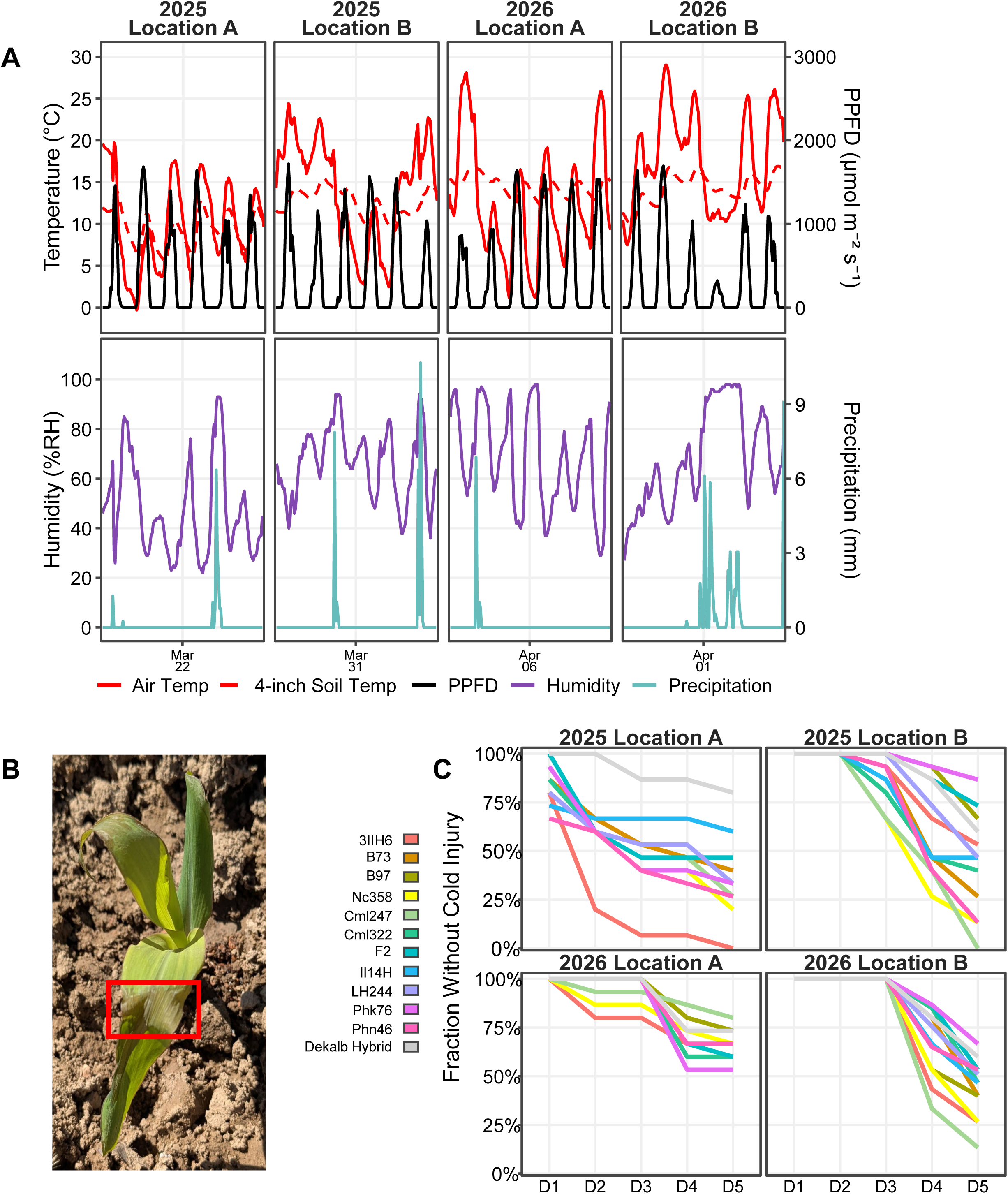
Environmental conditions and visible cold injury during four early-spring field experiments. A) Air temperature, 4-inch soil temperature, PPFD, relative humidity, and precipitation during the 2025 and 2026 experiments at Locations A and B. B) Representative plant from the 2025 Rollins; the red box identifies damaged leaf tissue. C) Resistance to cold injury, expressed as the fraction of plants within each genotype that had not developed visible cold injury on each day following transplanting. Lines show block-averaged genotype values for the four field environments. The experimental design comprised three replicate blocks per environment, with five plants per genotype per block.

### Field expression of genotypic variation is strongly environment dependent

Morphometric and physiological traits fluctuated markedly among field environments, and the expression of genotype effects was not uniform across sites or years (Fig. 6). Assimilation differed significantly among genotypes in both Location A environments but not in either Location B environment. In contrast, light-adapted F_V_′/F_M_′, immature-leaf growth, and relative mature-leaf SPAD showed significant genotype differences in both Location B environments but not in the corresponding Location A environments. Genotype rankings also shifted among environments, and no single genotype was consistently superior across traits although the commercial check variety (Dekalb Hybrid) was frequently among the top performers in all environments in which it was evaluated.

**Figure 6.**
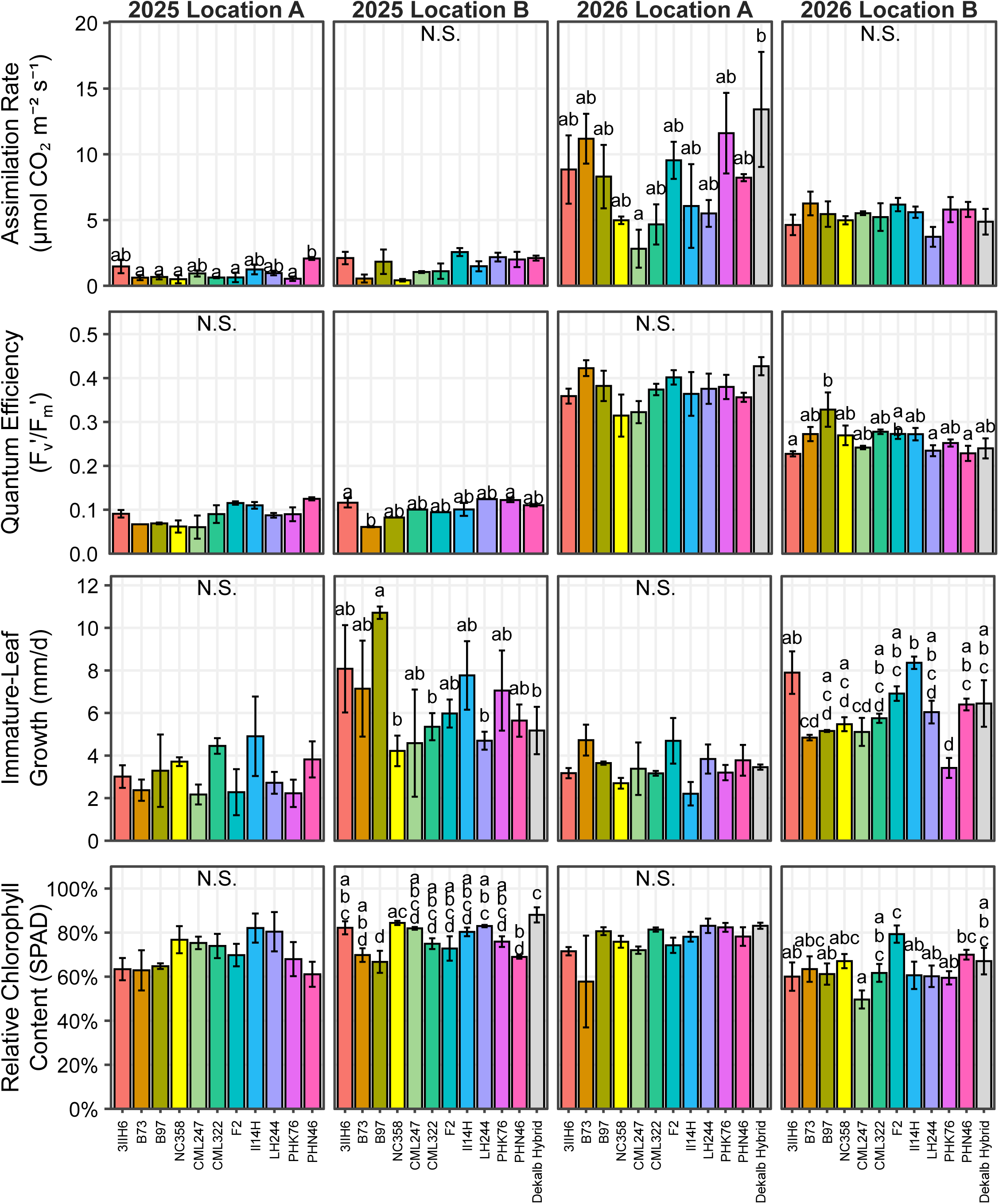
Morphometric and physiological variation among maize genotypes in four early-spring field environments. Columns show the 2025 and 2026 experiments at Locations A and B. Rows show net CO₂ assimilation at five days after transplanting, light-adapted Fv′/Fm′ measured on the same sampling date, immature-leaf growth over the five-day field exposure, and mature-leaf relative chlorophyll content (SPAD) at day 5 expressed relative to the pretransplant value. Values are means of block-level experimental units and error bars represent SE across replicate blocks. Experiments contained three replicate blocks, with five plants per genotype per block targeted; plant subsamples within a block were summarized to a single block-level value before statistical analysis. Different letters indicate Tukey-adjusted genotype comparisons at P < 0.05 within an environment; N.S. indicates a nonsignificant genotype effect. Individual-environment analyses used genotype as a fixed effect and block as a random effect.

Across all four field environments, assimilation increased with ΦPSII. A moderate positive correlation was observed among the pooled block-level dataset (Fig. 7). The distribution was strongly structured by environment, however, and environment-specific regressions ranged from little apparent relationship in 2025 Location A to much tighter positive relationships in the 2026 sites. Thus, ΦPSII remained positively associated with carbon assimilation across the combined multi-year multi-environment experiment, but the degree of association depended strongly on the environment in which plants were measured.

**Figure 7.**
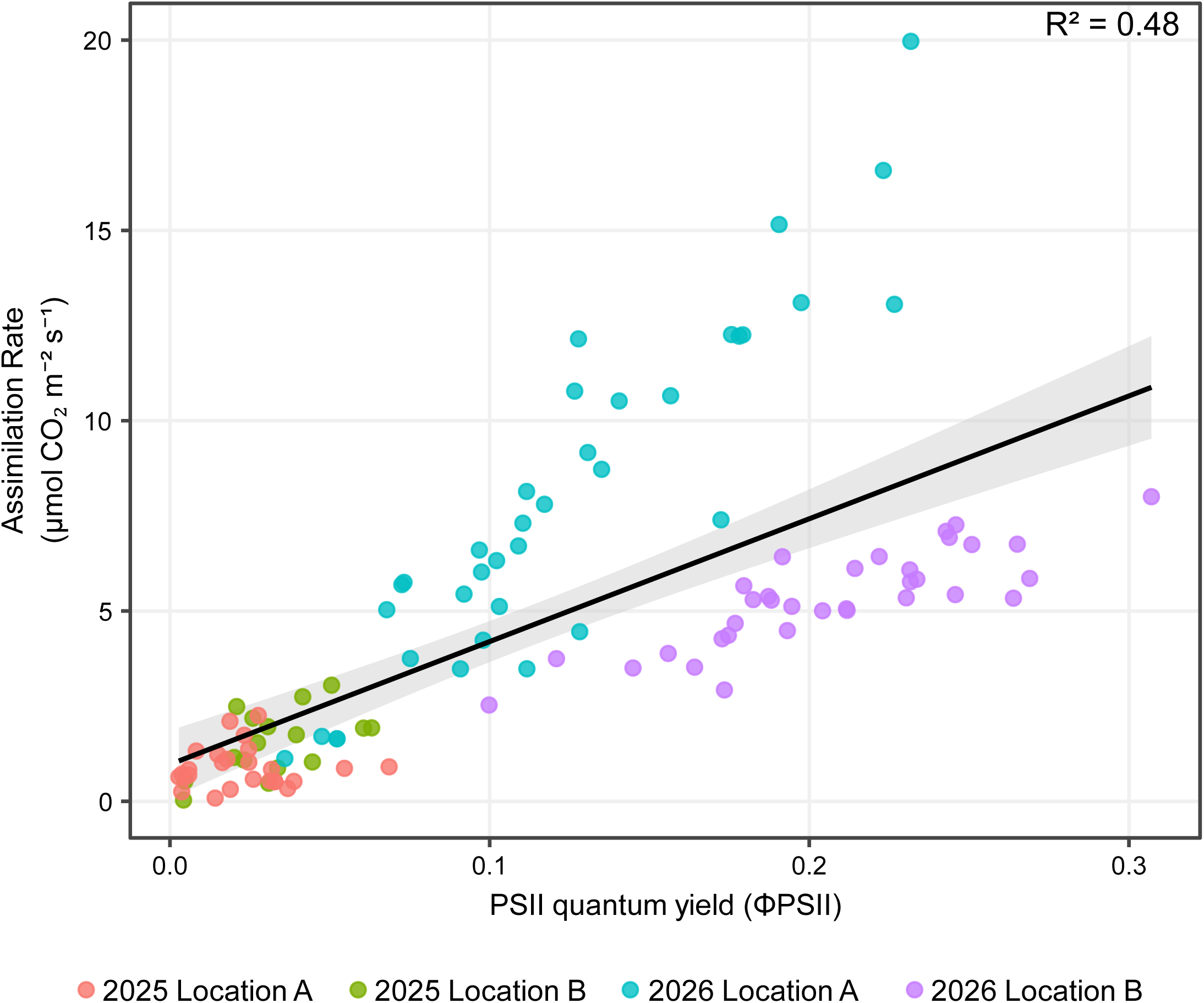
Relationship between net CO₂ assimilation and ΦPSII across four field environments. Net CO₂ assimilation is plotted against ΦPSII for block-level observations collected from the 2025 and 2026 field experiments at Locations A and B. Points are colored by field environment. The black line and shaded region show the pooled ordinary least-squares fit and its 95% confidence interval across all environments. The pooled relationship had an R2 = 0.48; N =112. The plotted regression is a descriptive ordinary least-squares fit to the block-level observations.

Additional field relationships were consistent with the broader physiological response. Greater endpoint assimilation was positively associated with greater immature-leaf growth accumulated over the same five-day exposure (Supplementary Fig. S10). By contrast, stomatal conductance showed no clear positive association with mature-leaf SPAD at the field endpoint (Supplementary Fig. S11). Together with the controlled-environment results, these field observations support maintenance of carbon assimilation and PSII operating efficiency as more closely coordinated components of early-season performance than chlorophyll status and stomatal conductance alone.

### Field traits differ substantially in broad-sense heritability

The cross-source validated panel of 1,310 SNPs was distributed across all 10 maize chromosomes and revealed substantial variation in realized genomic relatedness among the 11 inbred lines (Supplementary Fig. S2). Principal-component analysis separated the most genetically distinct genotypes and recapitulated broad germplasm relationships, with PC1 and PC2 accounting for 36.2% and 14.2% of marker variation, respectively (Supplementary Fig. S3).

Broad-sense heritability estimated across the four field environments differed markedly among traits (Table 1). In the primary phenotype-BLUP analysis, entry-mean H² was highest for mature-leaf SPAD both before field exposure (0.73) and after chilling exposure (0.86). ΦPSII showed moderate to high heritability (0.55). Assimilation was moderately heritable (0.45), whereas immature-leaf growth was lower (0.32) and light-adapted F_V_′/F_M_′ showed comparatively little heritable variation (0.22). The complementary GBLUP estimates reinforced the high heritability of SPAD and the low heritability of light-adapted F_V_′/F_M_′, while estimates for the remaining growth and photosynthetic traits shifted in magnitude after genomic relatedness was incorporated.

**Table 1.** Broad-sense heritability of maize morphological and photosynthetic traits across field environments. Heritability estimates were obtained from multi-environment mixed models fitted to genotype × environment × block means from 11 maize inbred lines evaluated across four field environments. Phenotype BLUP estimates treat genotype effects as independent and identically distributed and represent the primary phenotypic heritability analysis, whereas GBLUP estimates incorporate realized genomic relationships among the inbred panel. For the phenotype BLUP model, entry-mean H2 was calculated from estimated genotype, genotype × environment, and residual variance components for the four-environment, three-block design. Cullis H2 is a prediction-error-variance-based measure of the reliability of pairwise genotype differences, and mean ΔH2 is the mean of the corresponding pair-specific reliability estimates. For GBLUP, Cullis H2 and mean ΔH2 were calculated using the genomic variance expected for each pairwise genotype difference to account for unequal genomic relatedness among lines.

|  | **Phenotype BLUP** |  |  | **GBLUP** |  |
|---|---|---|---|---|---|
| *Trait* | *Entry-mean H²* | *Cullis H²* | *Mean Δ H²* | *Cullis H²* | *Mean Δ H²* |
| Mature leaf SPAD (prechill) | 0.73 | 0.73 | 0.73 | 0.83 | 0.78 |
| Mature leaf SPAD (after chill) | 0.86 | 0.86 | 0.86 | 0.92 | 0.89 |
| Immature leaf growth | 0.32 | 0.32 | 0.32 | 0.51 | 0.44 |
| Assimilation rate | 0.45 | 0.45 | 0.45 | 0.45 | 0.38 |
| ΦPSII | 0.55 | 0.51 | 0.51 | 0.40 | 0.33 |
| F’<sub>V</sub>/F’<sub>M</sub> (light adapted) | 0.22 | 0.20 | 0.20 | 0.00 | 0.00 |

Phenotype BLUP is the primary broad-sense heritability analysis; GBLUP is complementary. Δ H² is the difference-based reliability estimate.

Taken together, the controlled-environment and field experiments identify a common dynamic response: chilling rapidly restricts carbon assimilation, PSII operating efficiency, leaf elongation and developmental progression, with prolonged stress compressing the expression of genetic differences. Both photosynthetic function and development are restored when favorable temperatures return, revealing substantial variation in the extent and rate of recovery among maize genotypes. The persistence of an assimilation–ΦPSII relationship under controlled and field conditions, together with measurable field heritability for both traits, identifies photosynthetic maintenance and recovery as experimentally tractable components of the cold-tolerant early-planted maize ideotype. The V-stage and biomass allocation results further show that recovery includes renewed maturation of leaf tissue, linking leaf-level physiological recovery with whole-plant developmental progression.

## Discussion

### Maize chilling resilience is defined by both photosynthetic maintenance and recovery

The capacity of maize to experience a temporary reduction of photosynthetic capacity during chilling followed by its quick return with favorable temperatures represents complementary components of early-season cold resiliency. This contrasts with the concept of cold tolerance, which describes the ability of maize to maintain photosynthetic capacity throughout the chill stress. The results shown here depict resiliency over tolerance through declines of assimilation, ΦPSII, and stomatal conductance upon exposure to chilling, that rebounded to baseline level or greater when the chill was lifted. The drops seen were consistent with the established sensitivity of maize photosynthesis to temperatures below its optimum and indicate that the initial limitation to carbon gain was not attributable solely to stomatal closure (Long *et al*., 1983; Garzon *et al*., 2020). Convergence of physiology under severe limitation should not necessarily be interpreted as an absence of genetic variation for chilling resilience. Rather, prolonged exposure may drive diverse genotypes toward a common low-function state in which differences in physiological capacity become difficult to resolve, a model which has been previously postulated (Riva-Roveda *et al*., 2016).

The photosynthetic resiliency of maize to chilling is a complex phenotype encompassing multiple physiological processes and potential breeding targets (Burnett and Kromdijk, 2022). That concept was seen here where genotypes differed in the magnitude of recovery, demonstrating that restoration of photosynthetic function is itself a variable trait. These differences are biologically important because episodic chilling in agricultural environments is followed by periods in which plants must rapidly resume carbon gain and development. Moreover, chilling stress is known to sensitize the photosynthetic apparatus to light intensity (Nie *et al*., 1995; Szalai *et al*., 1996) and an early spring recovery period is likely to be accompanied by high incident irradiance, as evidenced by the field experiments reported here.

Photosynthetic recovery can occur at different rates among processes; classical studies of chilling-dependent photoinhibition in maize demonstrated that components of carbon assimilation and photochemical efficiency need not recover synchronously (Ortiz-Lopez *et al*., 1990; Fryer *et al*., 1995). Chilling injury is often characterized as tolerance by measurements made during or immediately following stress, whereas these results follow the recovery process of light-adapted fluorescence, dark-adapted F_V_/F_M_, and nonphotochemical energy dissipation immediately after the stress is lifted as well as five days later. The collective responses of those photosynthetic parameters show that the reduction and subsequent recovery of carbon assimilation are governed through coordinated changes across carbon metabolism, photochemistry, photoprotection, and ultimately photosystem function. This is consistent with previous work concluding that chilling sensitivity is not monocausal (Foyer *et al*., 2002; Long and Spence, 2013).

### Sustained chilling and high light disrupt coordination of carbon assimilation and photochemistry

In many controlled-environment studies of maize, chilling has been imposed with low to moderate irradiances (150–300 µmol m⁻² s⁻¹), in some cases specifically to limit photooxidative damage during cold exposure (Capell and Dörffling, 1993; Irigoyen *et al*., 1996; Szalai *et al*., 1997, 2018; Aroca *et al*., 2001, 2003; Kościelniak and Biesaga-Kościelniak, 2006). The use of sustained high irradiance (1,200 μmol m⁻² s⁻¹) comparable to field conditions captures an important physiological challenge relevant to clear spring days in the U.S. Corn Belt. That is, leaves must intercept substantial solar radiation while cool air and soil temperatures constrain the ability to use that energy.

The differing speed and magnitude of change in the response to chilling across the maize panel demonstrates that chilling did not affect photosynthesis through a single regulatory step, instead pointing towards nonstomatal limitation and a concurrent restriction of photochemical energy utilization. In general, operating PSII efficiency remained closely related to carbon assimilation even as absolute photosynthetic rates changed markedly among temperature stages and genotypes. Changes in NPQ were consistent with those of photoprotective energy dissipation as the capacity to use absorbed light declined. Reductions in dark-adapted F_V_/F_M_ after sustained chilling provide evidence that prolonged stress extended beyond the rapidly reversible regulation of operating efficiency to persistent changes in maximum PSII efficiency. These responses are consistent with the broader understanding of maize chilling injury as a consequence of imbalance among light capture, electron transport, carbon metabolism, antioxidant capacity, and repair of the photosynthetic apparatus (Fryer *et al*., 1995, 1998; Aroca *et al*., 2001; Foyer *et al*., 2002). Genotypic differences following the return to favorable temperatures may reflect both the severity of accumulated impairment and the capacity to restore functional coordination among carbon metabolism and photochemistry.

The present experiments cannot independently partition the effects of low temperature, irradiance, and their interaction because irradiance was not manipulated factorially during chilling. Instead, exposing maize to sustained chilling while maintaining substantial light availability created an integrated physiological environment relevant to the conditions encountered by seedlings planted earlier into temperate production systems. Future factorial experiments varying both temperature and irradiance could determine which genotypes are particularly sensitive to excess excitation pressure during chilling and whether genotypic differences in recovery originate primarily from metabolic limitation, accumulated photoinhibition, or their interaction.

The acclimation of maize to suboptimal temperatures has been well described in the literature; such acclimation processes have shown to contribute to subsequent tolerance of more extreme chilling. Leaves developed at 15 °C exhibited greater antioxidant capacity and photoprotective energy dissipation and sustained less photooxidative damage during successive exposure to 5 °C and high irradiance than leaves developed at 25 °C (Leipner *et al*., 1997), while growth at moderately low temperature induces broader adjustments in photosynthetic metabolism and antioxidant status (Kingston-Smith *et al*., 1999). Thus, the transition from favorable conditions to sustained chilling imposed here represents only one form of cold stress that may present in an early-spring field environment. Seedlings may experience gradual cooling, repeated cool nights, disproportionate chilling of the root zone, or shorter cold events interspersed with favorable conditions, each of which may alter the balance among hydraulic, metabolic, and photoprotective responses (Melkonian *et al*., 2004; Riva-Roveda *et al*., 2016). Consequently, genotypic performance under the present regime should be interpreted as tolerance to sustained chilling under high light rather than as a complete representation of low-temperature resilience. Importantly, however, field experiments indicated that the controlled-environment treatment captured physiologically relevant cold responses, with genotypic patterns observed in the growth chamber broadly translating to field performance despite the greater stochasticity and factors which are difficult to reproduce experimentally.

The C_4_ species *Miscanthus × giganteus* is known for its ability to productively function at temperature ranges below that of modern maize (Głowacka *et al*., 2014; Bilska-Kos *et al*., 2018). The *Miscanthus* accessions included here did not outperform maize across all measured traits, emphasizing that cold resilience is not represented by a universally superior value for any single fluorescence or gas exchange parameter. Nevertheless, the broader *Miscanthus* comparison remains an important physiological benchmark: productive C_4_ photosynthesis under cool conditions is biologically achievable, and improvement of maize will likely require coordinated changes across multiple components of photosynthetic function.

### Genetic variation in recovery translates across whole-plant and field responses

Variation in photosynthetic recovery was accompanied by differences in growth and whole-plant performance, indicating that the consequences of chilling extended beyond instantaneous leaf-level carbon exchange. Immature-leaf growth was strongly affected by chilling, consistent with temperature-dependent limitations of cell division and expansion (Ben-Haj-Salah and Tardieu, 1995; Rymen *et al*., 2007; Louarn *et al*., 2010), and growth resumed to differing degrees following return to favorable temperatures (Lainé *et al*., 2024; Ojeda-Rivera *et al*., 2025). Likewise, the restoration of assimilation in mature leaves does not guarantee the rapid restoration of whole-plant growth that is needed to overcome the stress induced limitation of new leaf area production and expansion.

Recovery of both leaf growth and development are critical to the chilling resiliency of maize to ensure minimal loss of future photosynthetic area. Renewed developmental progression can increase future carbon capture even when individual leaves do not fully return to their prechill physiological state. The combined gas-exchange and growth responses observed here therefore support a broader definition of chilling recovery that includes the restoration of physiological function and resumption of developmental activity, both of which impact biomass accumulation. Differences in biomass accumulation reflect the cumulative consequences of carbon acquisition, respiration, developmental rate, and allocation over time and therefore cannot be assigned to any single photosynthetic trait.

Chlorophyll related traits provide an additional, although more indirect, dimension of leaf-level responses to chilling and high light. SPAD exhibited substantial genetic repeatability and may therefore provide a comparatively tractable phenotype for large-scale screening. However, chlorophyll retention should not be interpreted as equivalent to maintenance of photosynthetic function. Leaves can retain chlorophyll while experiencing substantial reductions in carbon assimilation or operating photochemical efficiency, particularly during short-duration stress (Ting *et al*., 1991; Andrews *et al*., 1995). This creates an important practical distinction for breeding programs: traits that are highly heritable and inexpensive to measure are not necessarily the traits most closely aligned with the physiological process of interest. Additionally, the most mechanistically direct measurements may be difficult to deploy at scale.

### Physiological cold resilience provides a route toward earlier-planted maize

The agronomic importance of traits described here becomes clearest when chilling resilience is considered within the context of the maize planting window. Earlier planting has contributed to historical increases in maize productivity in the central United States by increasing the effective growing season and facilitating the use of germplasm adapted for a longer duration growing season. Across 12 central U.S. states from 1979 to 2005, it was estimated that trends toward earlier planting for half of the states accounted for 19–53% of state-level yield increases (Kucharik, 2008). Additional calendar days afforded by earlier planting only contribute to productivity if plants can develop photosynthetically active leaf area and maintain growth under the chill stresses encountered during that period (Bollero *et al*., 1996). Thus, genotypes capable of maintaining greater photosynthetic activity during moderate chilling, limiting accumulated photochemical impairment under high light, and rapidly restoring carbon gain and leaf development following stress are needed to make early-season plantings successful.

The results presented here should not be taken as a counterfactual to existing crop modeling assumptions; these experiments do not directly establish how much grain yield would be gained by improving assimilation, ΦPSII, or photosynthetic recovery during chilling. Instead, the present results identify genetic variation in physiological processes that could contribute to improving grain yield of early-season maize plantings. This is particularly relevant to the CERCA cold-tolerant maize ideotype, in which early-season cold resilience is an enabling trait rather than an endpoint (Ojeda-Rivera *et al*., 2025). Photosynthetic cold resilience directly links the seedling stress phenotypes to broader agronomic objectives: maintenance of carbon assimilation during chilling to preserve immediate carbon gain; maintenance of ΦPSII and effective photoprotection to help sustain productive use of intercepted radiation; rapid recovery of carbon acquisition following transient stress; and quick restoration of leaf development to increasing photosynthetic photon capture.

In total, these results demonstrate substantial standing variation in the capacity of maize to maintain and recover photosynthetic function and growth following chilling under high irradiance. Carbon assimilation, operating PSII efficiency, photoprotection, maximum PSII efficiency, leaf development, and whole-plant growth respond on different timescales and differ in their degrees of genetic variation. This complexity argues against defining chilling resilience by a single endpoint or trait, and it provides multiple physiological routes for crop improvement. Of these, maintenance and recovery of carbon assimilation and ΦPSII provide a particularly direct connection between the leaf-level response to chilling and the ability of an early-planted crop to productively use captured solar radiation. Describing the heritable genetic variation of photosynthetic maintenance, photochemical regulation, recovery, and development provides a route toward that objective. In this way, the physiological variation identified here establishes a foundation for developing maize capable of converting an earlier planting window into greater seasonal carbon assimilation, more effective resource acquisition, and ultimately a more productive and resource efficient crop.

## Acknowledgements

We thank Harper LaFond (USDA-ARS) and Amy Litherland (University of Missouri) for their assistance with logistics and data collection for all experiments. We thank Ray Wright (USDA-ARS) and Chris Browne (University of Missouri) for their aid with the field experiments.

## Author Contributions

TJM, ENL, JDW: Conceptualization; TJM, ENL, JDW: methodology; TJM, ENL: formal analysis; TJM, ENL: investigation; JDW: resources; TJM and ENL: writing—original draft; TJM, ENL, DC, JDW: writing—review & editing; TJM and JDW: data curation; TJM: visualization; JDW: funding acquisition.

## Funding

This research was supported by the United States Department of Agriculture (USDA) Agricultural Research Service (USDA-ARS), including funding from projects 8062-21000-052-005-T. TJM and DC were supported by ORISE postdoctoral fellowships, ENL was supported by an ORISE predoctoral fellowship.

## Data Availability

The primary data supporting this study were not made publicly available at the time of publication. The data that support the findings of this study are available from the corresponding author upon request

## Supplementary Data

**Supplementary Figure S1.**
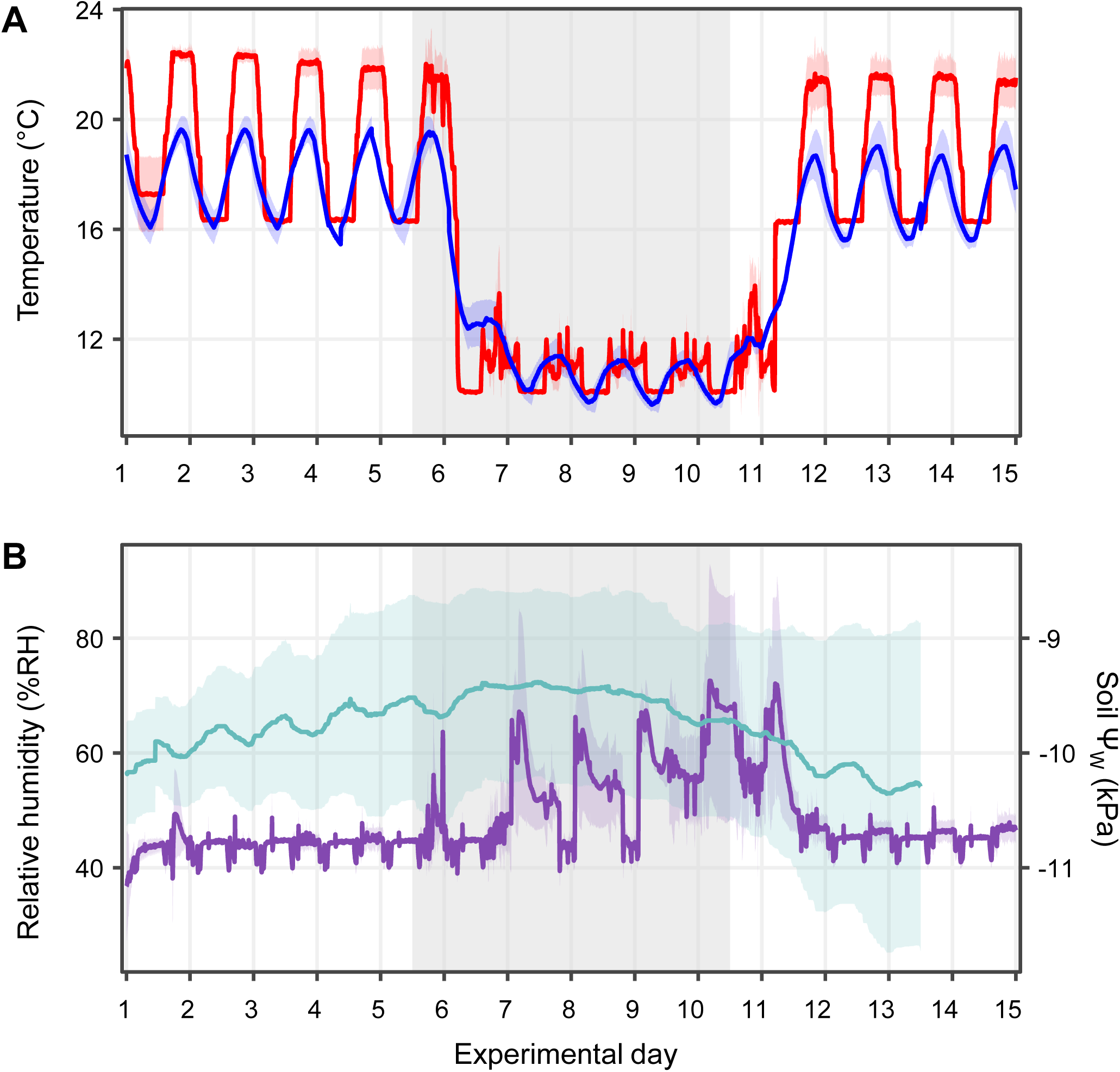
Environmental conditions imposed during controlled-environment experiments. (A) Air and soil temperature across the 15-day experimental sequence comprising five days of prechilling, five days of chilling, and five days of recovery. (B) Relative humidity and soil water potential (fi_W_) over the same experimental sequence. The shaded interval denotes the chilling treatment. Environmental records were compiled from four experiments conducted under the common treatment regime; lines denote mean values and shaded ribbons display 95% confidence interval.

Fig. S2. Genome-wide distribution of SNP markers and resulting genomic relationship matrix

**Supplementary Figure S3.**
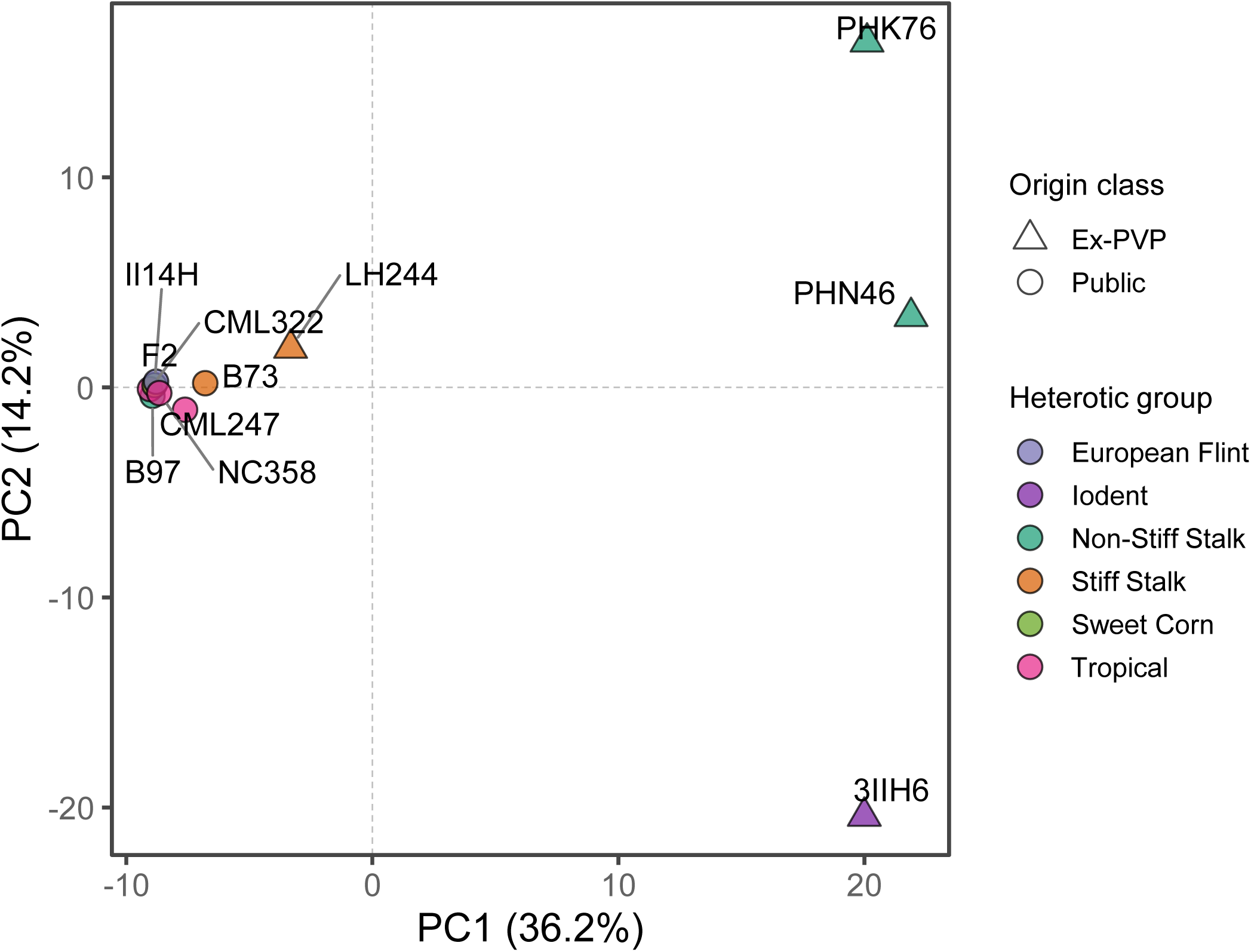
Population structure of the maize inbred panel based on the validated marker set. Principal component analysis of the 11 maize inbred lines using the same Platinum marker matrix used to construct the genomic relationship matrix. PC1 and PC2 account for 36.2% and 14.2% of marker variation, respectively. Point colors identifies germplasm/heterotic group, while the outer symbols denote origin (Public or Ex-PVP).

**Supplementary Figure S4.**
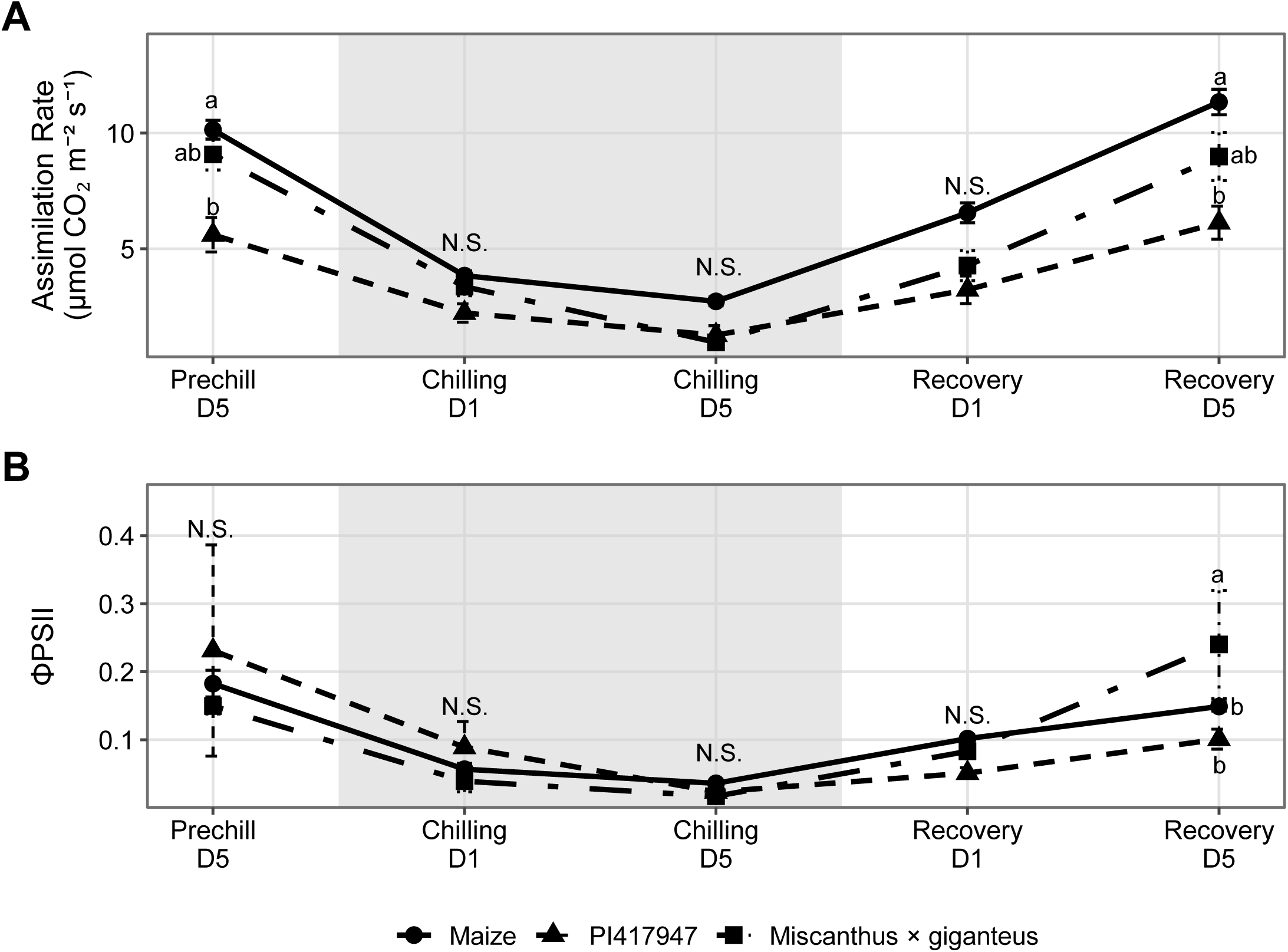
Photosynthetic physiology of maize and two *Miscanthus* accessions across prechilling, chilling, and recovery. (A) Net CO₂ assimilation and (B) ΦPSII are shown for the pooled maize panel, *Miscanthus* accession Pi417947, and *Miscanthus × giganteus* at five experimental timepoints. The shaded interval denotes the chilling treatment. Multiple instrumental measurements from the same plant and timepoint were averaged before analysis, and each trait was restricted to experiments providing complete five-timepoint support for both maize and *Miscanthus*. Points are model-estimated marginal means ± SE. N= 56 maize plants and 7-10 miscanthus replicates. Letter denote comparison of the three groups within a timepoint; groups not sharing a letter differed at Tukey-adjusted P < 0.05. Statistical inference was made using repeated measures models which included group × timepoint and experiment × timepoint effects, as well as a genotype effect.

**Supplementary Figure S5.**
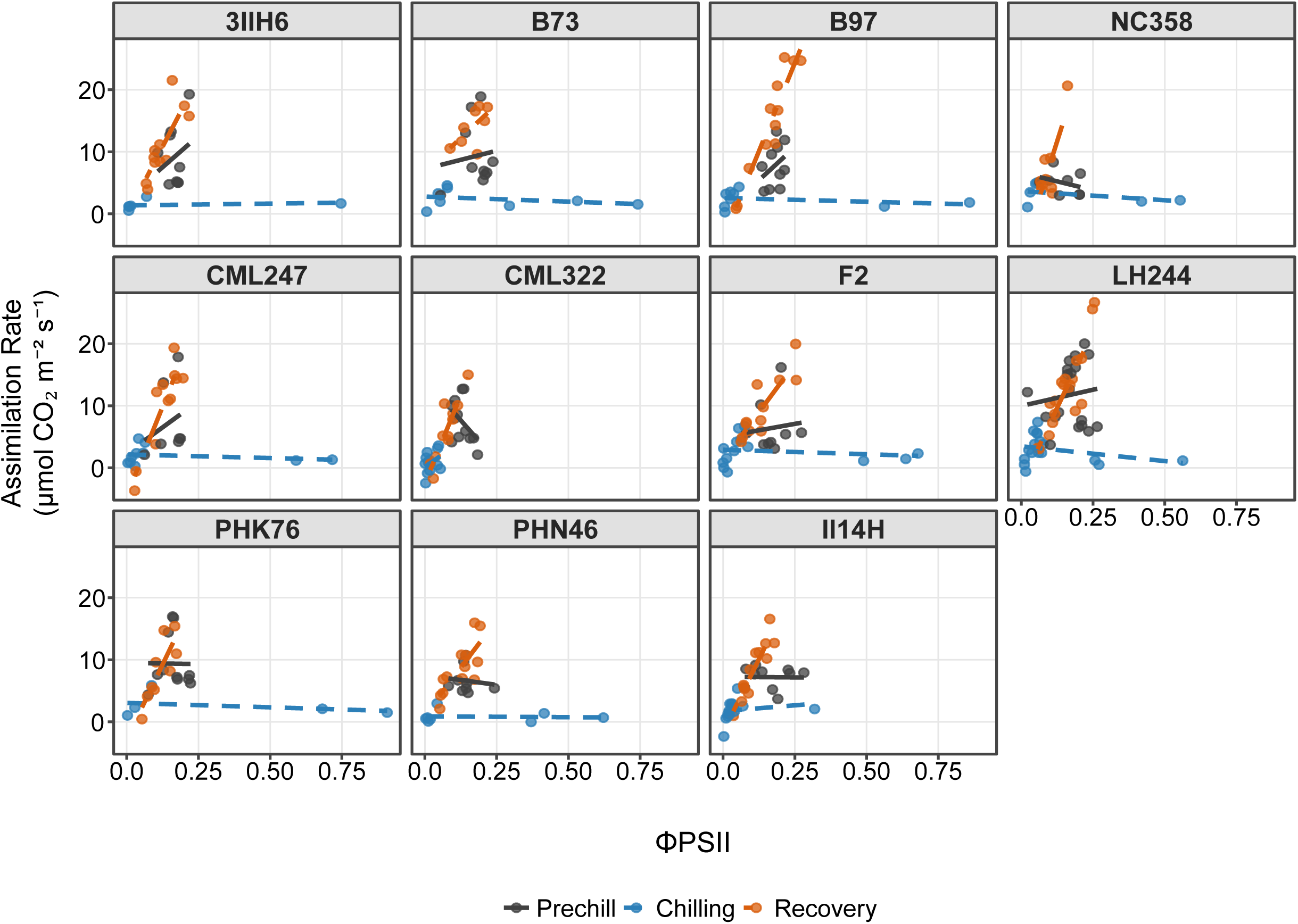
Effect of genotype on the relationship between net CO₂ assimilation and ΦPSII across experimental stages. Net CO₂ assimilation is plotted against ΦPSII for each of the 11 maize genotypes at Prechill D5, Chilling D5, and Recovery D5. Points represent paired individual biological observations, and fitted lines show genotype and stage specific least-squares. *N* = 114 Prechill, 111 Chilling, and 128 Recovery observations (*N* = 353 total). Fitted lines are descriptive genotype × stage regressions.

**Supplementary Figure S6.**
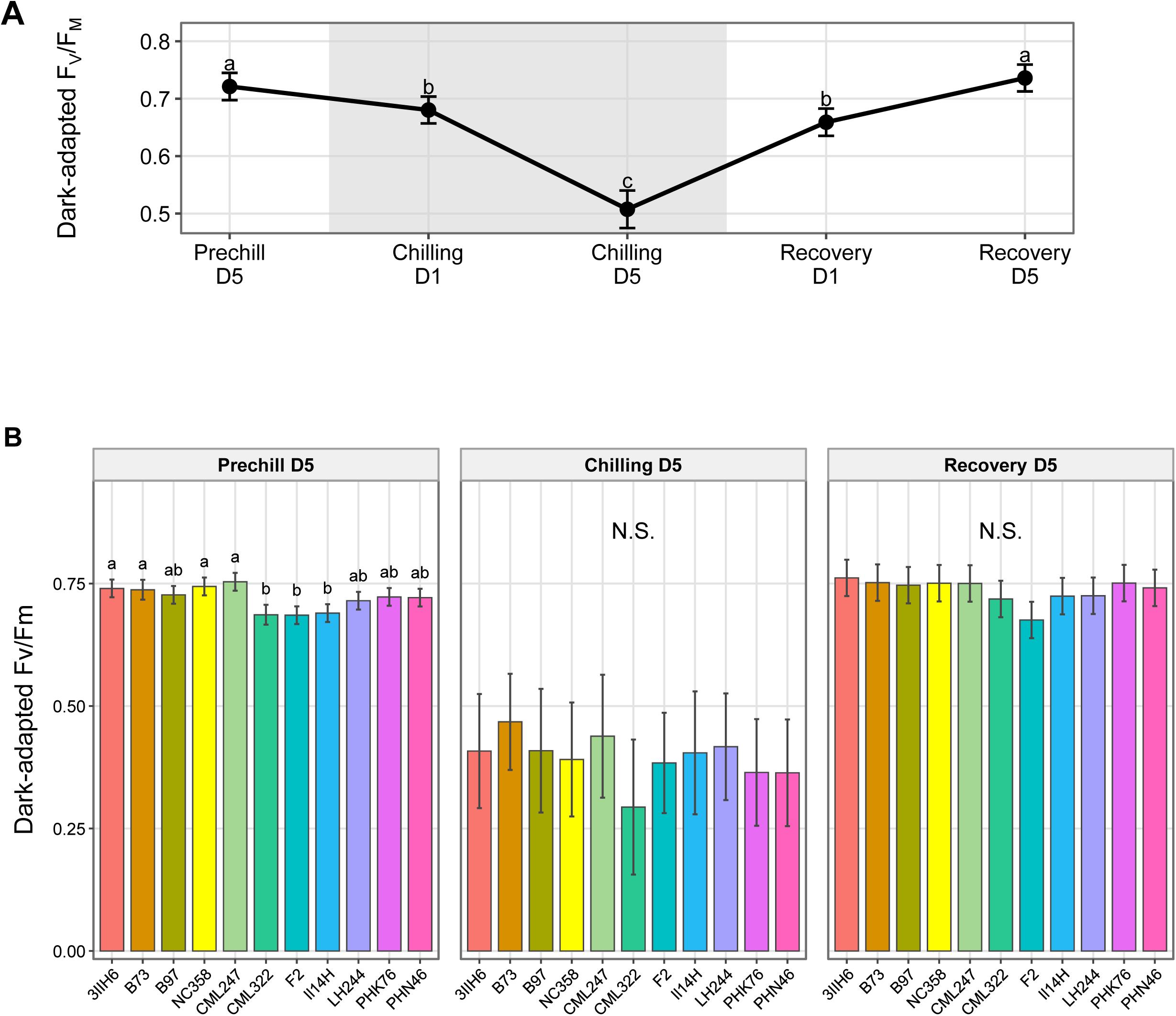
Dark-adapted Fv/Fm across the chilling treatment and among maize genotypes. (A) Model-adjusted dark-adapted Fv/Fm across the course of the experiment; the shaded interval denotes the chilling treatment. Points are estimated marginal means with associated SE shown with error bars. (B) Dark-adapted Fv/Fm for each of the 11 maize genotypes at Prechill D5, Chilling D5, and Recovery D5. Bars/points represent genotype EMMs with associated 95% confidence intervals shown by error bars. N= 67. In (A), different letters indicate significant differences among timepoints; in (B), letters compare genotypes within each stage, with groups not sharing a letter differing at Tukey-adjusted *P* < 0.05; N.S. indicates *P* ≥ 0.05.

**Supplementary Figure S7.**
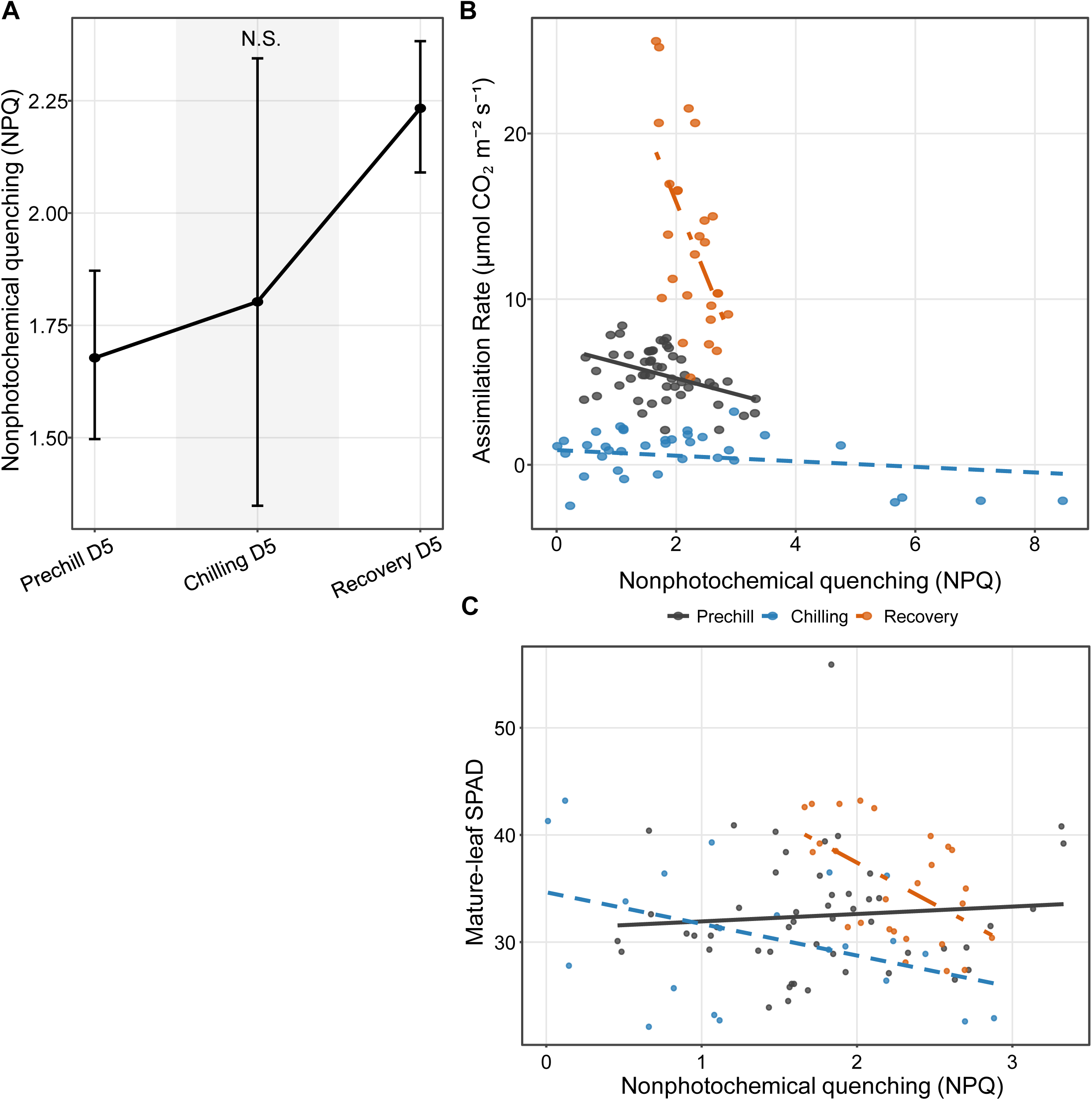
Nonphotochemical quenching across experimental stages and its relationships with carbon assimilation and mature-leaf chlorophyll status. (A) Nonphotochemical quenching (NPǪ) at Prechill D5, Chilling D5, and Recovery D5. Points and error bars represent estimated marginal means and 95% confidence intervals from stage-specific models; the shaded interval denotes the chilling stage. (B) Relationship between net CO₂ assimilation and NPǪ at the D5 endpoint of each experimental stage. (C) Relationship between mature-leaf SPAD and NPǪ at the same stage endpoints. In (B) and (C), points represent individual plants with matched measurements, lines show stage-specific linear relationships. N = 51, 36, and 26 NPǪ observations at Prechill D5, Chilling D5, and Recovery D5, respectively.

**Supplementary Figure S8.**
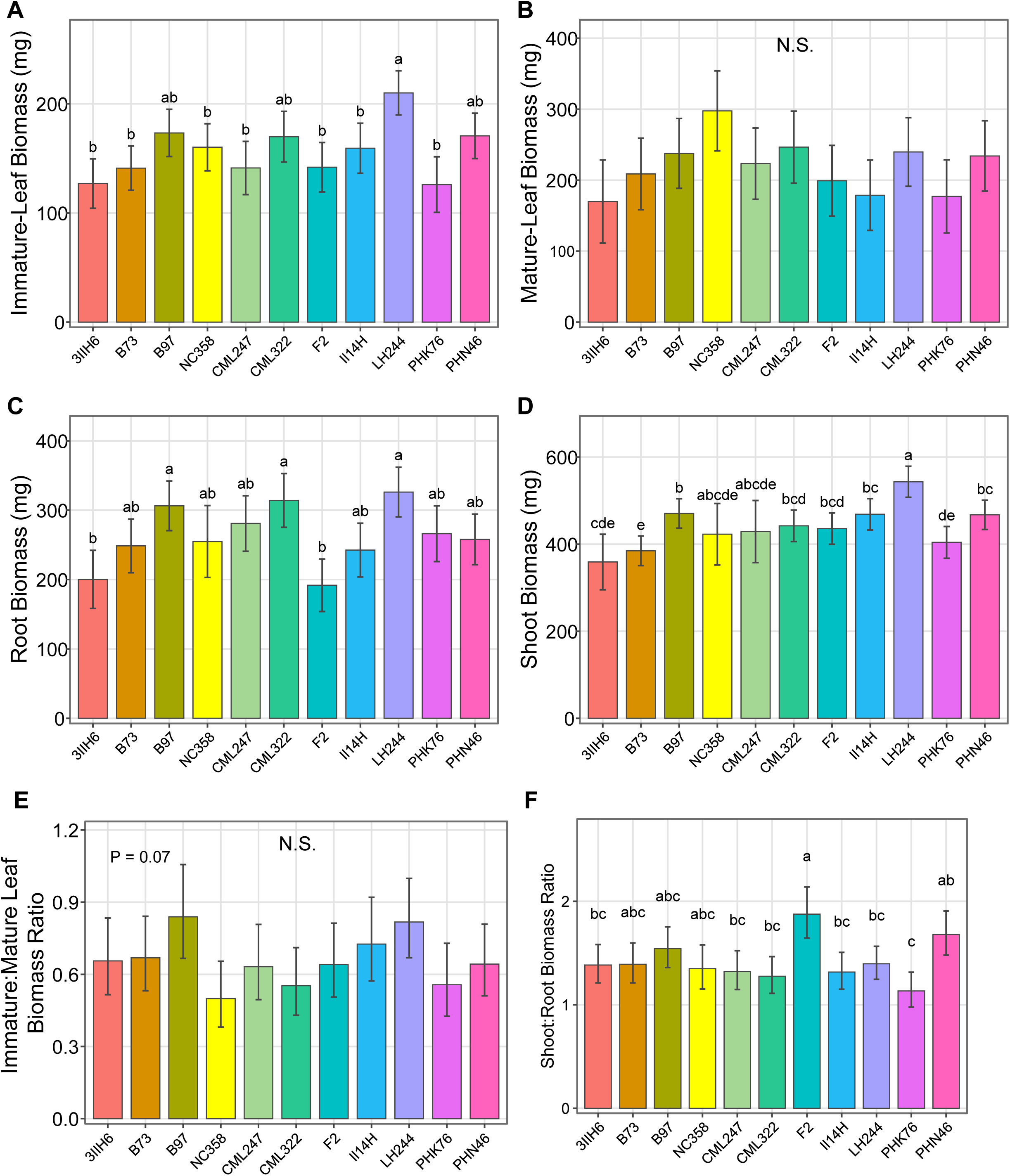
Dry weights of (A) immature leaves, (B) mature leaves, (C) roots, and (D) shoots (mature leaves, immature leaves, and stem tissue), E) Immature:mature leaf biomass ratio. F) Shoot:root biomass ratio measured at harvest. Values are genotype EMMs with 95% CI displayed as error bars; biomass ratios were analyzed on the log scale and are displayed as ordinary biomass ratios. N= 195

**Supplementary Figure S9.**
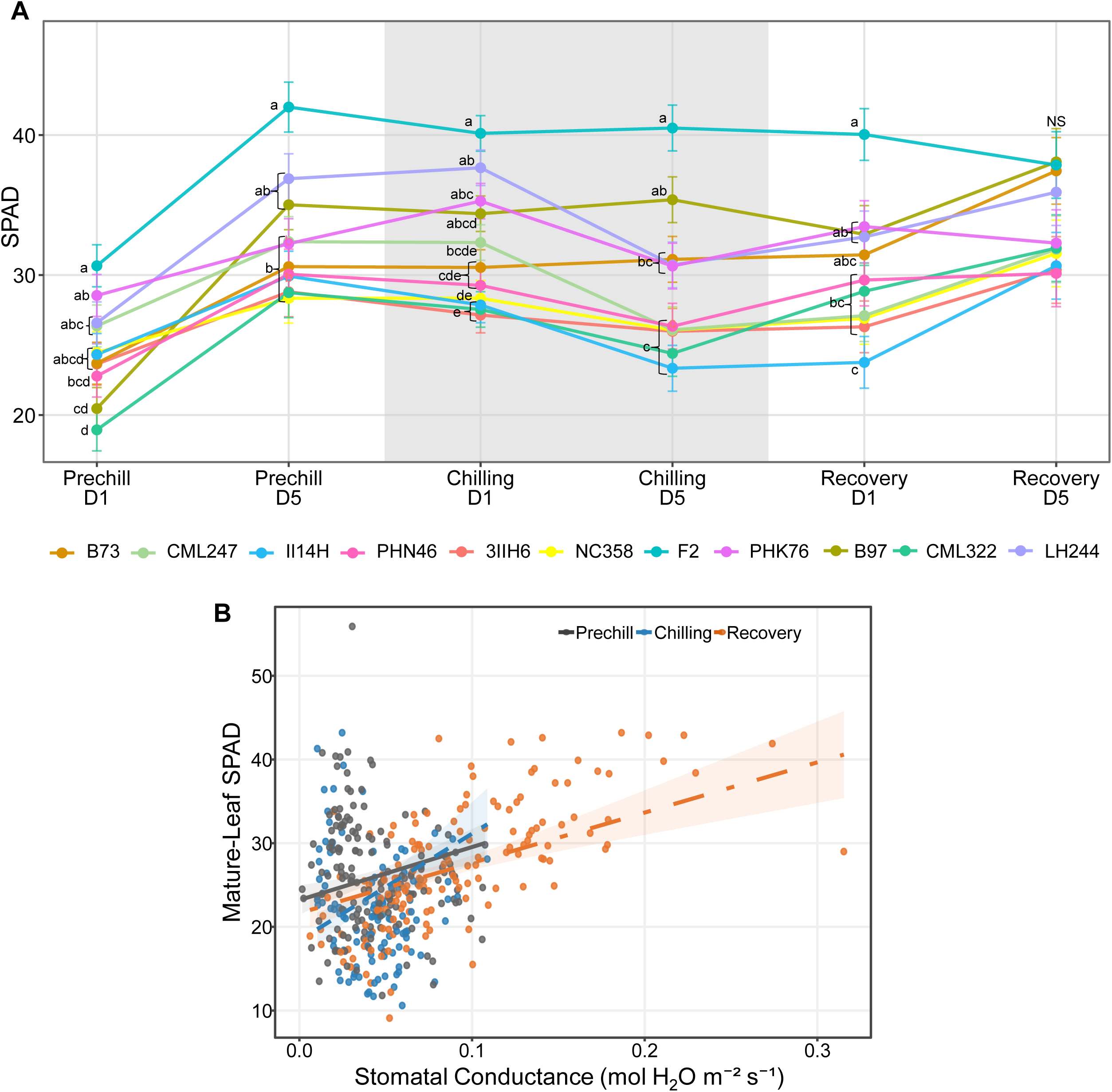
A) Genotypic variation in mature-leaf chlorophyll content across prechilling, chilling, and recovery. Mature-leaf relative chlorophyll content (SPAD) is shown for each maize genotype at Prechill D1 and D5, Chilling D1 and D5, and Recovery D1 and D5. The shaded interval denotes the chilling treatment. Points are genotype EMMs and error bars represent SE. N = 66 plants. Within each timepoint, genotypes not sharing a letter differed at Tukey-adjusted P < 0.05; N.S. indicates that the genotype effect was not significant P *<* 0.05. B) Mature-leaf SPAD is plotted against stomatal conductance (g_sw_) at the D5 endpoint of Prechill, Chilling, and Recovery. Points represent individual plants with matched SPAD and gas-exchange measurements. Lines show the stage-specific linear relationships; shaded bands indicate 95% confidence intervals. N = 153, 117, and 148 matched observations for Prechill, Chilling, and Recovery, respectively. Stage-specific linear models included g_sw_ and replicate experiment as fixed effects; g_sw_ was significantly associated with SPAD within each stage (P<0.05).

**Supplementary Figure S10.**
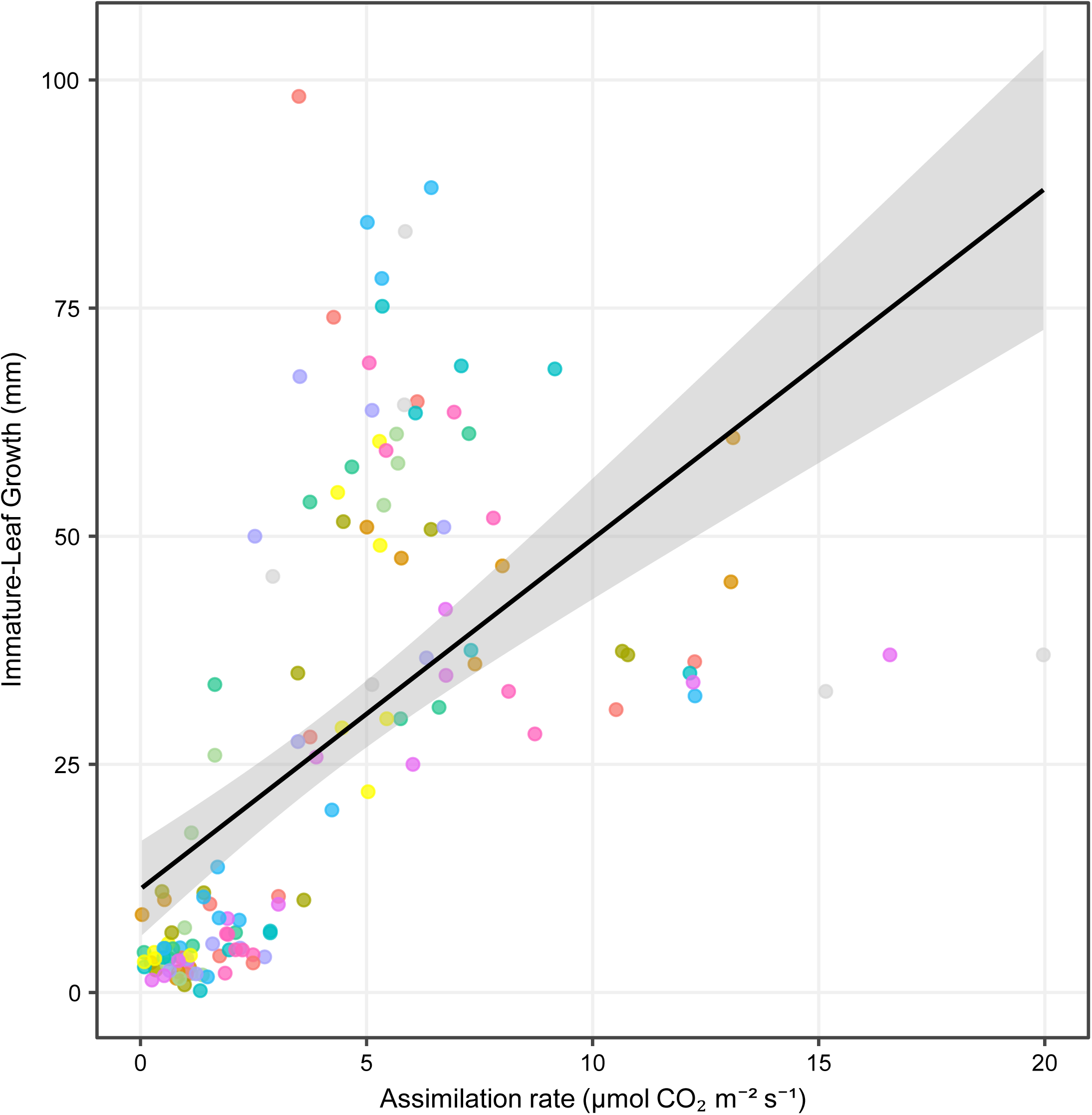
Relationship between net CO₂ assimilation and immature-leaf growth during field exposure. Immature-leaf growth over the five-day field interval is plotted against net CO₂ assimilation measured at five days after transplanting. Points represent block-level observations from the four field environments and are colored by genotype. The black line and shaded region show the pooled ordinary least-squares fit and 95% confidence interval. N = 135. The plotted line is an ordinary least-squares regression across the available block-level field observations.

**Supplementary Figure S11.**
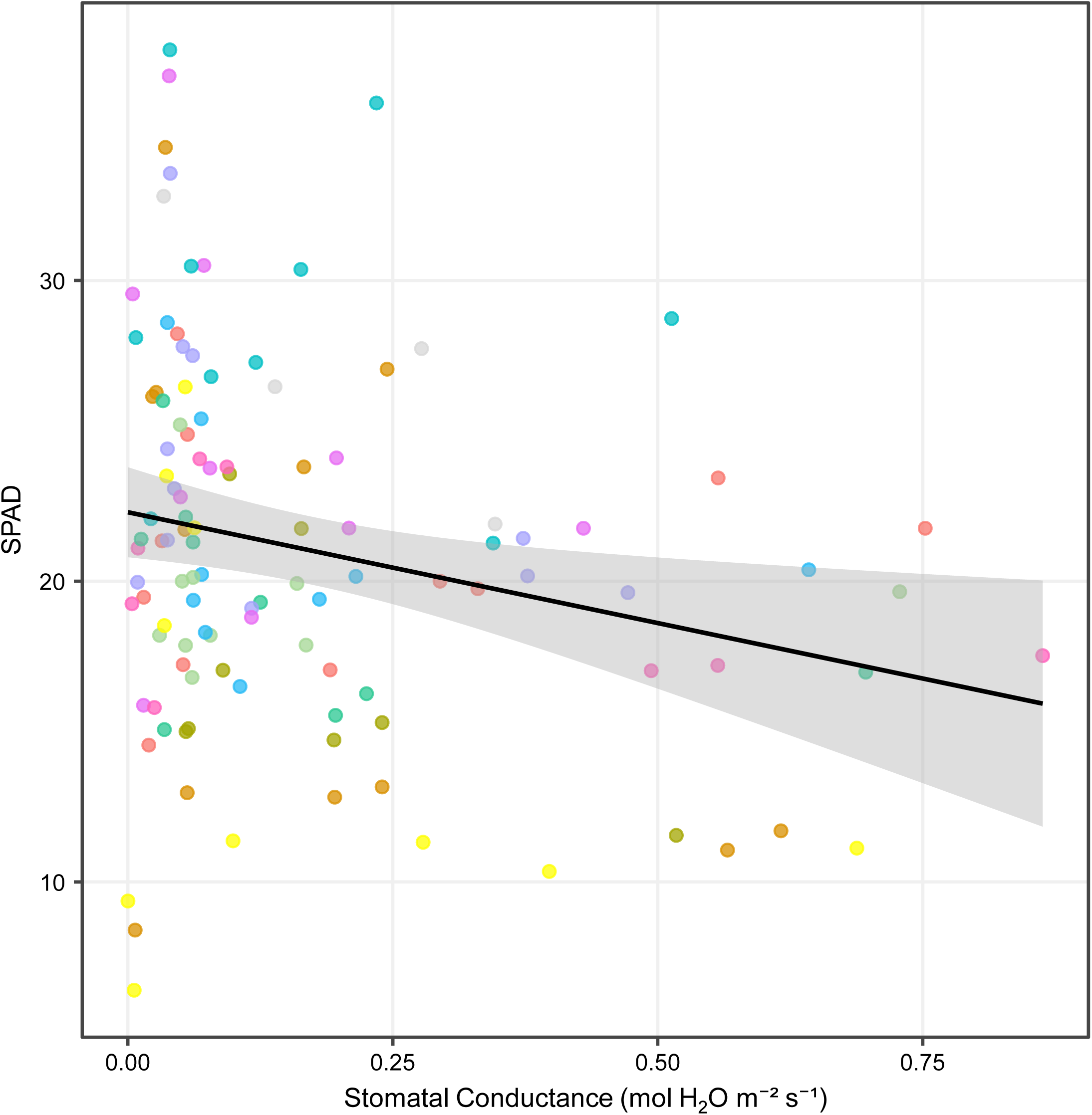
Relationship between stomatal conductance and mature-leaf chlorophyll content during field exposure. Mature-leaf SPAD measured at the end of the field exposure is plotted against stomatal conductance measured at the same endpoint. Points represent block-level observations from the four field environments and are colored by genotype. The black line and shaded region show the pooled ordinary least-squares fit and 95% confidence interval. N = 112. The displayed relationship is an ordinary least-squares regression across the available block-level field observations.

Supplementary Table 1. Model selection criteria

## Notes

### Competing Interest Statement

The authors have declared no competing interest.

